# Control of motility and cell shape of *Haloferax volcanii* is linked by a transcriptional regulator

**DOI:** 10.64898/2026.02.03.703544

**Authors:** Phillip Nußbaum, Felix Grünberger, Felix Neuschütz, Kevin Chou, Shamphavi Sivabalasarma, Alexander Eulitz, Anna-Lena Sailer, Katharina Vogl, Marten Exterkate, Wei He, Anita Marchfelder, Dina Grohmann, Sonja-Verena Albers

## Abstract

Archaea rely on motility and morphological plasticity to navigate their environments, yet the transcriptional regulation of these processes remains poorly understood. In *Haloferax volcanii*, archaellum-dependent motility is transcriptionally regulated, but an EarA-like master regulator is absent. Here, we identify CsmR as a transcriptional regulator that links archaellum biogenesis and cell-shape transitions in *H. volcanii*. Deletion of *csmR* abolished detectable motility, whereas overexpression increased motility and promoted a sustained rod-like morphology. Comparative transcriptomics defined a CsmR-associated regulon that includes archaellum and chemotaxis genes as well as rod-shape determinants (e.g., Sph3 and RdfA), and upstream motif enrichment supports a direct role for CsmR in transcriptional control. Furthermore, *csmR* and *cirA*, a KaiC-like regulator, share extensive transcriptional overlap, with CirA likely fine-tuning CsmR-mediated regulation through post-translational modification. These findings establish CsmR as a key integrator of motility and cell shape regulation in *Haloferax volcanii*, suggesting that haloarchaea coordinate these fundamental processes through an unidentified transcriptional network. Moreover, Northern blotting and cell shape observation suggest that transcription factor RosR is involved in the regulation of an sRNA that shares extensive overlap with the *cirA* gene, possibly fine-tuning the effect of CirA on the regulation of the archaellum cluster and the rod shape determinants *sph3* and *rdfA*. Understanding this interplay provides new insights into archaeal adaptability and may reveal broader regulatory principles in prokaryotic cell biology.

## Introduction

The ability of microorganisms for cellular locomotion is a key feature to evade unfavorable environmental conditions such as nutrient limitation, changes in pH or temperature. The most widespread form of cellular movement is swimming motility, in which cells use a rotating appendage to generate force and propel themselves through liquid media. Depending on the phylogenetic group, different cellular machineries have evolved, including cilia in Eukaryotes, flagella in Bacteria, and archaella in Archaea [1]. The archaellum shares homology with archaeal and bacterial type-IV pili with respect to the proteins involved and in their mode of assembly [2,3]. Across archaeal phyla, the archaellum is composed of different sets of proteins that share a conserved core machinery [4]. Those core components include the polytopic membrane protein ArlJ [5], the assembly ATPase ArlI [6], the ATP binding protein ArlH [7,8], stator proteins ArlF/ArlG [9] and the filament forming archaellins [10]. Typically, all archaellum-related genes are clustered together in a single operon [4,10,11]. In contrast to the bacterial flagellum, which is powered by the proton motive force [12], archaellum rotation is driven by ATP hydrolysis via ArlI [6,13]. Since ATP is essential for many cellular processes, this energy-intensive mode of locomotion must be tightly controlled. In Thermoproteota diverse transcriptional factors that control the expression of archaellum components have been described. In the thermophilic archaeon *Sulfolobus acidocaldarius*, several positive and negative regulators have been studied, showing that phosphorylation plays a major role in archaellum regulation [4]. The key components of this system collectively referred as archaellum regulatory network (Arn) include the Forkhead associated (FHA) -domain containing protein ArnA and a von Willebrand factor A domain containing protein called ArnB [14]. Both proteins undergo phosphorylation at their C-terminus by the kinase ArnC [14,15]. Additionally, ArnB is phosphorylated by a second kinase, ArnD [15]. Phosphorylation of ArnA and ArnB promotes their interaction and the formation of a higher oligomeric complex [16]. This complex negatively affects archaellum gene expression [14], possibly by binding the *arlB* or *arlX* promotor region, as observed for the ArnA homolog of *Sulfolobus tokodai*, ST0829 [17]. Other components of the archaellum regulatory network include the paralogous transmembrane proteins ArnR and ArnR1, which contain helix-turn-helix motifs in their N-terminal region [18]. The previously mentioned kinase ArnC and a third membrane-associated kinase ArnS have been shown to phosphorylate ArnR and ArnR1 [18,19]. Phosphorylation of both proteins had a positive effect on the activation of the *arlB* promotor, but the second promotor of the archaellum operon, upstream of *arlX*, remained unaffected by ArnR/R1 [18,20]. High upregulation of the archaellum cluster in *S. acidocaldarius* is seen in the late stationary phase of growth or under starvation conditions [21].

In Euryarchaeota a single transcription regulator of the archaellum cluster has been identified so far, named euryarchaeal archaellum regulator A (EarA) [22]. EarA was first characterized in the methanogenic archaeon *Methanococcus maripaludis* where deletion led to non-archaellated cells due to significantly reduced expression of one of the filament-forming subunits of the archaellum, *arlB2*. Electrophoretic mobility shift assays showed that EarA binds, via its winged helix-turn-helix motif, to a region upstream of the archaellum promoter [23]. Interestingly, a spontaneous mutation in the B recognition element (BRE) of the *arlB2* promoter restored archaellation and motility in the *earA* deletion strain. These BRE mutations generated a strong promoter motif, probably bypassing the need for EarA to recruit the transcription factor B (TFB) to the archaellum promoter [24]. These findings suggest that EarA acts as a positive regulator for archaellum expression. Indeed, *earA* overexpression in *Pyrococcus furiosus* resulted in heavily archaellated cells [25].

While EarA has been established as the primary archaellum regulator in many Euryarchaeota, its absence in Haloarchaeota suggests that motility regulation in this group follows a distinct molecular mechanism [22]. Instead of relying on a single dedicated activator, archaellation in halophilic archaea appears to be influenced by environmental factors, cell shape transitions, and potentially alternative regulatory networks. In *Haloarcula marismortui* the salt concentration of the medium influences the composition of the archaellum filament [26], while in *Haloferax volcanii* lower growth temperatures induced strong upregulation of archaellins A1 [27]. Additionally, the ability of *H. volcanii* to swim is closely linked to its cell shape. During early exponential growth *H. volcanii* cells are rod-shaped and motile. As growth progresses, the cells transition from this rod-to a plate-shaped cell morphology in which cells are not motile [28,29]. Plate shaped cells lack archaellum filaments but retain assembled archaellum motors [28]. The formation of rod-shaped *H. volcanii* cells is controlled by several factors. One of those factors is CetZ1, a tubulin homolog that is essential for the formation and maintenance of motile rod-shaped *H. volcanii* cells [30]. A recent study identified three additional proteins involved in the morphological plasticity of *H. volcanii*. Rod-Determination Factor A (RdfA) is essential for the rod-shaped morphology, as its deletion leads to predominantly disk-shaped cells. Conversely, Disk-Determining Factor A (DdfA) and Sph3 are critical for maintaining a disk-shaped morphology, with its deletion resulting in exclusively rod-shaped cells. The third protein, Volactin, an actin homolog, contributes to disk formation through dynamic polymerization and depolymerization [31]. Moreover, a KaiC homolog in *H volcanii* named CirA was shown to impact cell shape and motility. The deletion strain of *cirA* stayed rod-shaped during all growth stages resulting in increased motility compared to the wildtype [32].

Here, we identify CsmR (<u>C</u>ell <u>s</u>hape and <u>m</u>otility regulator) as a novel transcriptional regulator of archaellation in *H. volcanii*. Through genetic, transcriptomic, and structural analyses, we demonstrate that CsmR is essential for motility and additionally plays a critical role in coordinating cell shape transitions. Unlike previously described archaellum regulators, CsmR belongs to the Lrp/AsnC family of transcription factors, a widely distributed regulator group known for integrating environmental and metabolic signals. Our findings reveal that CsmR directly regulates archaellin expression and likely acts in the same regulatory network as CirA, providing a new link between motility, cell morphology, and environmental adaptation in archaea. These discoveries offer fresh insight into archaeal regulatory networks and establish a new paradigm for archaellum control beyond EarA-dependent systems in Euryarchaeota.

## Results

### CsmR is a novel potential regulator of archaellation in *H. volcanii*, distinct from EarA

The gene encoding CsmR (*hvo_1209*), a small 13 kDa protein, is located upstream of the archaellin genes *arlA1* and *arlA2* in the *H. volcanii* genome. The presence of a predicted winged-helix DNA-binding domain suggested that similar to the previously identified positive regulators EarA in *M. maripaludis* and *P. furiosus* [22,25], this protein would be involved in the regulation of the archaellum cluster expression in *H. volcanii*. However, homology searches revealed no sequence or structural similarity between CsmR and EarA, suggesting a distinct regulatory mechanism in *H. volcanii*. Structural modeling classified CsmR within the Lrp/AsnC family of transcriptional regulators, known for their roles in metabolic and stress-response pathways in archaea and bacteria [33,34]. The highest prediction of confidence of the AlphaFold 3 model supported dimer formation (pTM 0.79), consistent with the dimeric nature of Lrp/AsnC regulators. The predicted dimerization interface of CsmR is mediated by a β-sandwich fold, further aligning it with Lrp/AsnC regulators rather than the previously characterized archaellum regulators (S1 Fig).

### Deletion of *csmR* results in a complete loss of motility whilst overexpression renders cells to be hypermotile

To investigate the function of the potential archaellum regulator in *Haloferax volcanii* we generated a marker less deletion strain of *csmR* and assessed a potential effect on cell locomotion via motility plates. In comparison to the control strain H26 the *csmR* deletion strain fully lost the ability to swim on semi solid agar plates (Fig 1A) indicating that *csmR* controls archaellum expression. Indeed, transmission electron microscopy of a Δ*pilB3*Δ*csmR* strain showed cells completely devoid of archaellum filaments (Fig 1B), whilst the Δ*pilB3* control strain showed archaellation. Since the discrimination between pili and archaellum filaments due to their homology is difficult, a deletion strain of the pilin assembly ATPase PilB3 [35] of *H. volcanii* was used to obtain cells that only have archaella as cell appendages.

**Fig 1.**
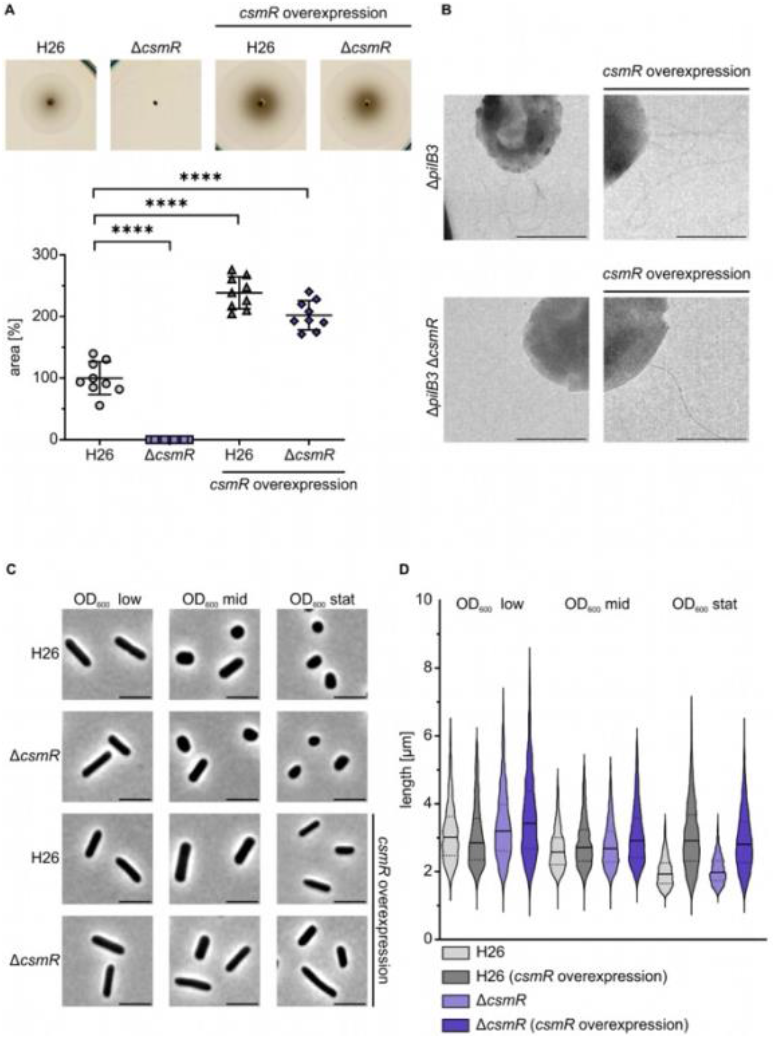
Deletion of *csmR* leads to a complete loss of motility whilst overproduction results in hypermotility. (A) Motility assay comparing wildtype H26 with the *csmR* deletion strain and both strains overexpressing *csmR*. Exemplary motility halos of all strains are shown. The area of the motility halos was measured and normalized to the average area of wildtype halos showing a significant reduction (p≤0.0001) in cell locomotion of the *csmR* deletion strain compared to H26 and a significant (p≤0.0001) increase of motility by *csmR* overproduction. Samples were measured in biological and technical triplicates and all nine single data points per strain were plotted. The middle line indicates the mean and the upper and lower line the standard deviation. (B) Transmission electron microscopy images of the *pilB3* and the *pilB3 csmR* double deletion strain (left) and of both strains overexpressing *csmR* (right). The filament of the archaellum is not assembled in the double deletion strain whilst overexpression of *csmR* resulted in assembly of an archaellum filament in the double deletion strain. Scale bar 1 µm. (C) Cell shape analysis of the wildtype H26 and the *csmR* deletion strain at low, mid and stationary OD_600_ and under *csmR* overexpression condition. Both strains show a transition from rod-to plate-shaped cells. In contrast both overexpression strains stay rod-shaped over the whole growth cycle and do not transition from rod-to plate shaped cells. Scale bar 4 µm. (D) Cell shape was analyzed using MicrobeJ and the summarized results of three independent biological replicates per strain and OD_600_ value plotted as violin-plots. The median is indicated by the middle black line, dotted lines indicate the first and third quartile. For each condition more than 1000 cells were analyzed.

Several studies showed that motility of *H. volcanii* strictly depends on its cell shape [36]. *Haloferax volcanii* is pleomorphic being rod shaped and motile in lag- and very early exponential growth phases whilst during mid exponential growth phase cells undergo a morphologic transition to plate shaped cells that are not motile anymore even though the archaellum motor is still present [28]. To exclude the option that the *csmR* deletion strain is immotile due to cell shape effects, we imaged cells of the deletion strain at different growth stages and compared them to strain H26 (early (OD_600_ of 0.02), mid (OD_600_ of 0.2), and stationary growth (OD_600_ of 2.0)). Both strains showed the typical cell shape of *H. volcanii* during growth, with rod-shaped cells at early growth stages and a transition to plate-shaped cells when they reach the stationary phase (Fig 1C, 1D). To see if those morphological changes impact the composition of membrane lipid types the abundance of several lipid species present in *H. volcanii* was assessed during different growth phases (S2 Fig). Overall, only small changes were observed, which predominantly involve the lower abundant lipid species PA (phosphatidic acid), PE (phosphatidylethanolamine), MGD (monogalactosyldiacylglycerol), as well as the cardiolipins. The only major abundant lipid species that seems to be mildly influenced by the growth stage is PG. This might be explained by the fact that PG serves as a substrate for all cardiolipins, which in turn seem to be slightly upregulated at later growth stages, which is a common phenomenon observed in prokaryotes. Next, analysis of the lipid tail distribution of the four most abundant lipid species in *H. volcanii* was performed (S3 Fig). C40:0 (i.e., two times a phytanyl chain) is the predominantly observed form, which is in accordance with the literature [37]. Other lipid tail configurations do exist, in which both the level of desaturation as well as the carbon chain length can differ. In general, there seems to be a trend that with increasing optical densities, there is a decrease in saturated lipid tails, except for PG. Again, this might be explained by the fact that unsaturated PG serves a substrate for cardiolipins, which is more prominently produced during later growth stages. Altogether though, the overall changes in lipid composition (headgroup as well as tail configuration) are minor. Hence, it is unlikely that the observed changes in cell morphology during growth are caused by an altered lipid composition.

Overexpression of the euryarchaeal archaellum regulator (EarA) in other Euryarchaeota led to drastically increased archaellum formation [25]. To test if overexpression of CsmR in *H. volcanii* has a similar effect, H26 and the *csmR* deletion strain were transformed with a plasmid that harbored *csmR* under the control of a strong xylose inducible promoter [38]. Overexpression of *csmR* in the Δ*csmR* strain complemented the non-motile phenotype of the deletion strain not only back to wildtype levels but caused hypermotility (Fig 1A). The same was observed when *csmR* was overexpressed in H26, causing motility halos more than twice the size of the motility halo of the control strain. Increased cellular protein concentration of EarA in *Pyrococcus furiosus* cells lead to hyperarchaellation [25]. However, in contrast to *P. furiosus*, Δ*pilB3 H. volcanii* cells transformed with the *csmR* overexpression plasmid showed no hyperarchaellation phenotype. The Δ*pilB3*Δ*csmR* double deletion strain assembled archaellum filaments upon *csmR* overexpression, as observed with the transmission electron microscope (Fig 1B); however, the number of archaella was comparable to the control strains with or without *csmR* overexpression plasmid (Fig 1B).

As mentioned before, cell motility of *H. volcanii* strongly depends on its cell shape. Whilst the control strain H26 and the *csmR* deletion strain transition from rod to a disk-like shape during growth, both strains maintain a rod-shaped appearance upon *csmR* overexpression (Fig 1C, 1D). Therefore, CsmR not only seems to affect archaellum expression but possibly also regulates genes that are involved in rod formation and/or maintenance.

### *CsmR* deletion induces widespread upregulation of archaellum and chemotaxis genes

To further investigate how CsmR regulates motility and cell shape transitions, we analyzed its impact at the transcriptional level. We performed RNA-sequencing on both wild type and Δ*csmR* strains under three distinct growth conditions to: (1) assess the overall transcriptional effect of *csmR* deletion and uncover potentially associated regulatory networks and (2) determine whether these effects vary under conditions where cells typically lack appendages and adopt a disk-like morphology (Fig 2A). Principal component analysis revealed that the growth phases of the harvested cultures were the primary driver of transcriptomic variation, while the deletion of *csmR* had only a smaller impact. This supports the hypothesis that CsmR functions as a dedicated regulator of archaellation rather than on a global scale (Fig 2B). To further investigate gene-specific effects, we performed differential gene expression analysis across the growth phases (early (OD_600_ of 0.02), mid (OD_600_ of 0.2), and stationary growth (OD_600_ of 2.0)). In early exponential growth, Δ*csmR* cells showed significant upregulation of shape-relevant genes, including *sph3* (*hvo_2175*) and r*dfA* (*hvo_2174*), as well as chemotaxis-associated genes (Fig 2C). Despite the complete loss of motility, RNA-seq analysis revealed a significant upregulation of archaellum genes in Δ*csmR* (Fig 2C). *H. volcanii* features a unique gene arrangement, where archaellation and chemotaxis genes are interspersed and transcribed in different orientations. Despite this organization, Δ*csmR* exhibited consistent upregulation across entire transcriptional units, including ArlA1/A2, ArlC/E/F/G/G/I/J/CheF2, CheB/A/R/F1, CheY/C/D, and ArlD/HVO_1202 (Fig 2D). This contrasts with the expected results, given the strain’s impaired swimming ability, and deviates from regulatory patterns observed in other euryarchaeal archaellum regulators [24,25]. Taken together, the transcriptomic analysis implies that CsmR might act as a repressor.

**Fig 2.**
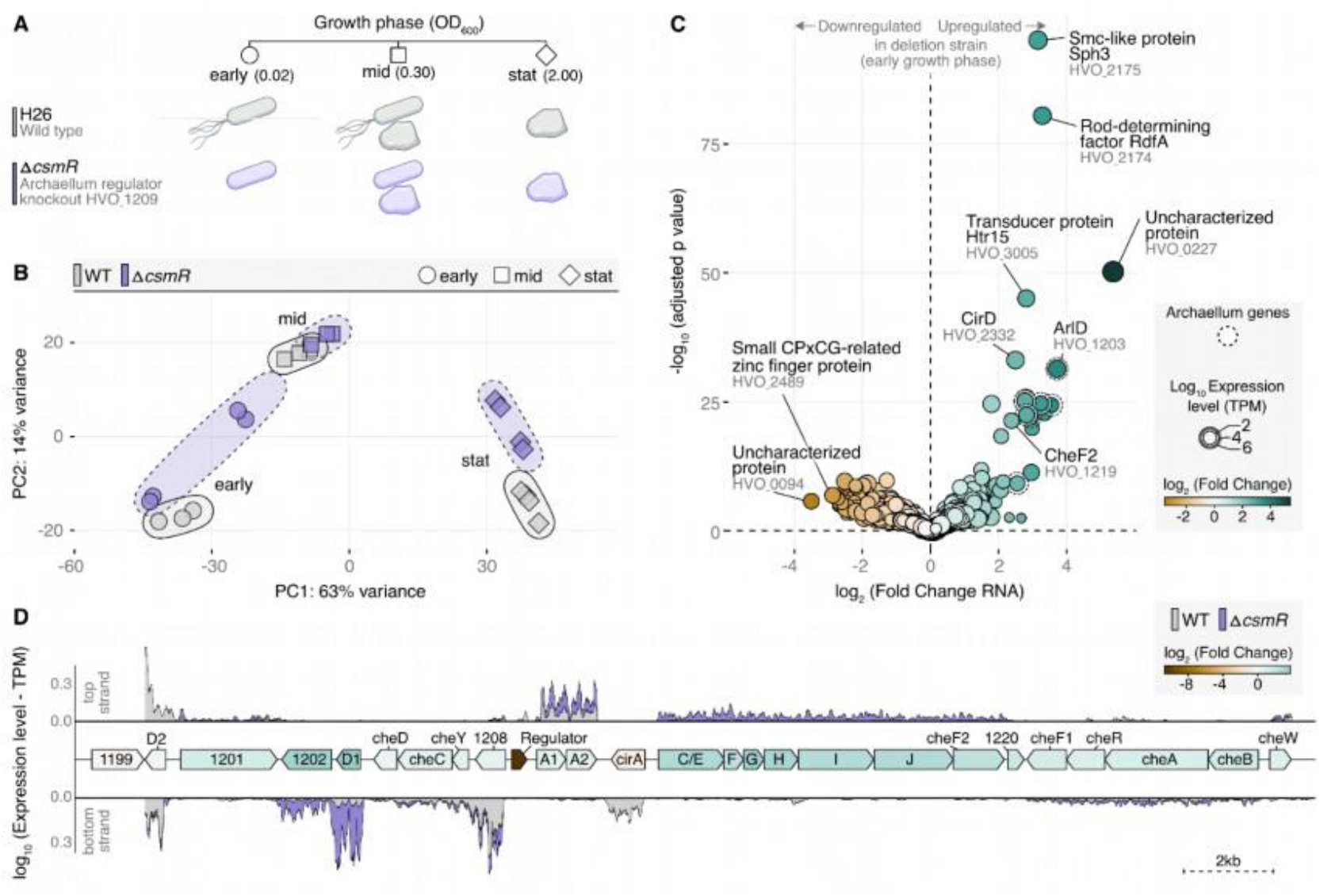
Deletion of *csmR* leads to upregulation of archaellum and chemotaxis genes. (A) Schematic of the experimental design, comparing the *Haloferax volcanii* wild type strain H26 and a deletion strain of the potential archaellum regulator CsmR, across different growth phases. Note that during normal growth conditions, *Haloferax* exhibits pleiomorphism, transitioning from rod-shaped, motile cells to sessile disk-like cells. (B) Principal component analysis from wild type (WT, grey) and Δ*csmR* (archaellum regulator, purple) strains, based on total read counts after outlier removal. Different growth phases are indicated by different cell shapes, illustrating the primary source of transcriptomic variance. (C) Volcano plot showing differential gene expression in low OD samples. Genes are represented as circles, with log_2_-fold changes color-coded from dark brown (downregulated in deletion strain) to dark green (upregulated in deletion strain), expression levels indicated by circle size and archaellum-related genes highlighted with dashed outlines. (D) Mean coverages of strand-specific, TPM (transcripts per million)-normalized reads are shown for wild type (grey) and *csmR* deletion strain (purple). The middle tracks display archaellum and chemotaxis-related genes, according to their relative gene size (size bar depicted). Genes are color-coded based on log_2_-fold changes from dark brown (downregulated in deletion strain) to dark green (upregulated in deletion strain) under low-OD conditions and strand orientation indicated by rectangle direction.

### CsmR deletion leads to persistent activation of motility and signalling pathways across growth phases

To assess the impact of the *csmR* deletion across different growth conditions, we next examined differential gene expression in mid and stationary samples. Pearson’s correlation analysis (R = 0.27) revealed a weak positive correlation between log_2_-fold changes in early and exponential (mid) growth phases, indicating that only a subset of genes was consistently up- or downregulated across both conditions (Fig 3A).

**Fig 3.**
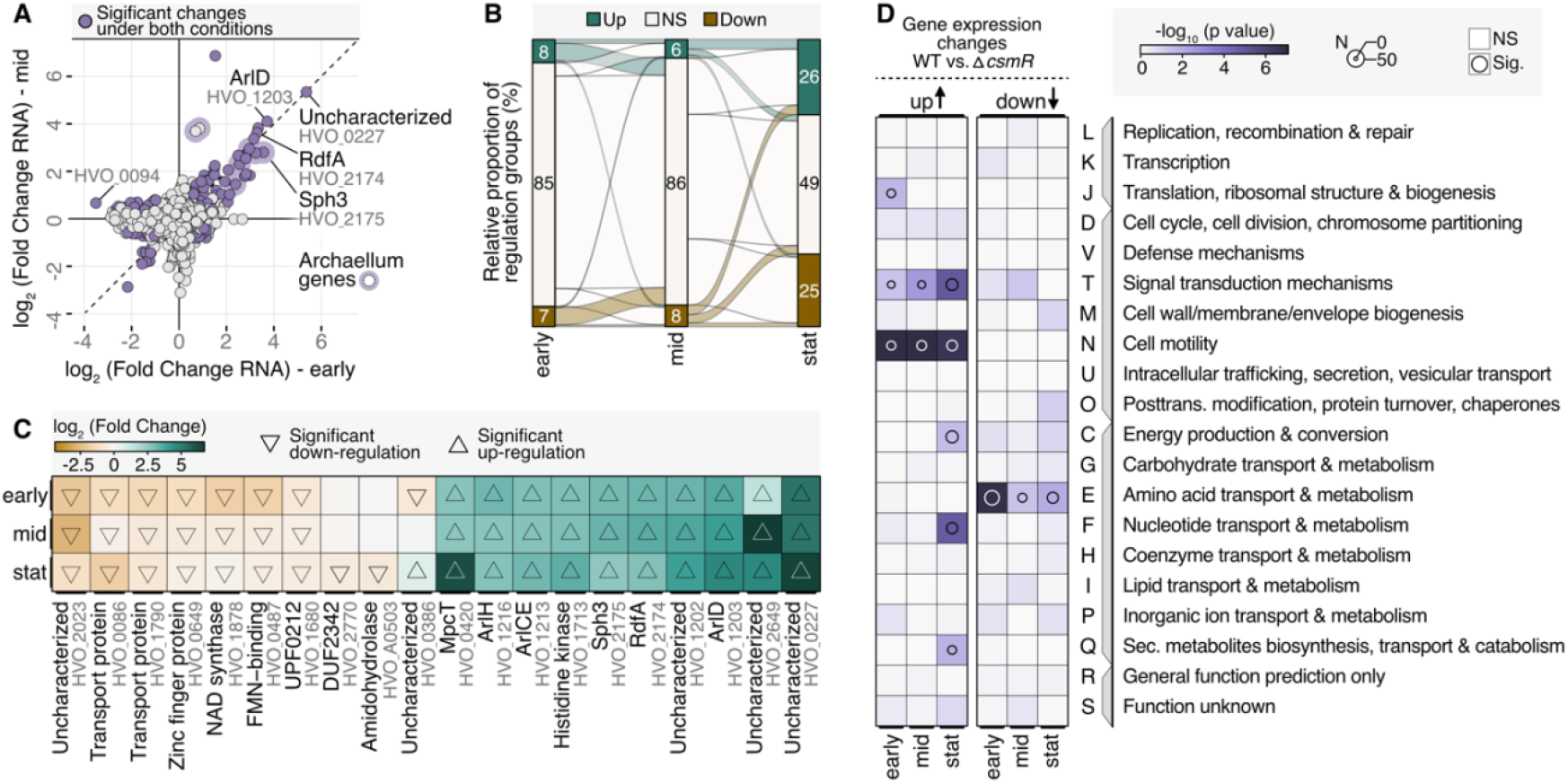
C*smR* deletion leads to a shared functional response across growth conditions. (A) Scatter plot showing the log_2_-fold changes in gene expression between early and mid-exponential samples for the Δ*csmR* strain relative to the wild type. Archaellum genes (dashed outline, light pink) and other genes that are consistently regulated under both conditions are highlighted. Genes with significant changes (adjusted *P*-value < 0.05) under both conditions are colored in pink. (B) Flow diagram illustrating the relative number and interconnectedness of genes between all growth conditions (x-axis) for each regulatory group (upregulated: green, downregulated: brown, rest: off-white). (C) Heatmap displaying the top 10 genes with the highest mean log_2_-fold changes across all conditions. Significance (adjusted *P*-value < 0.05) is indicated by triangles, with log_2_-fold changes color-coded from dark brown (downregulated in deletion strain) to dark green (upregulated in deletion strain). (D) Gene set enrichment analysis of archaeal clusters of orthologous groups (arCOGs) across all three growth conditions. Significantly overrepresented genes (*P*-value < 0.05, color-coded by a white to dark purple gradient) are marked by circles, with circle size representing the total number of differentially regulated genes detected in each group.

Notably, several of the most upregulated genes in early growth, particularly those linked to archaellation and cell shape regulation, remained upregulated in the exponential phase. Overall, during early phase, 8% of genes were upregulated and 7% downregulated, with similar proportions in the exponential phase (6% upregulated, 8 % downregulated), though there was minimal overlap between these gene sets (Fig 3B). In the stationary phase, where cells adopt a different morphology, gene regulation was significantly more pronounced, with 26% of the genes upregulated and 25% downregulated. Notably, archaellation and chemotaxis-associated genes remained consistently upregulated across all growth phases (Fig 3C). However, no clear functional pattern among downregulated genes could be detected. To further investigate the biological significance of these transcriptional changes, we performed functional enrichment analysis using archaeal clusters of orthologous groups (arCOGs). This analysis revealed significant overrepresentation in groups N (cell motility) and T (signal transduction mechanisms) among upregulated genes across all growth phases. Additionally, categories Q (secondary metabolites biosynthesis, transport, & catabolism), F (nucleotide transport & metabolism), and C (energy production & conversion) were enriched in stationary phase, while category E (amino acid transport & metabolism) was consistently downregulated across all conditions (Fig 3D).

### A conserved upstream motif is located upstream of genes of the CsmR regulon

To further interpret our findings - particularly the consistent upregulation of specific genes, suggesting shared regulation by CsmR - we performed motif detection analysis. We analyzed upstream sequences of genes that were significantly upregulated across all three growth conditions (adjusted *P*-value < 0.1, n: 62).

In 21 of these genes, we identified a semi-palindromic TATCA(N_4_)TGATA motif located approximately 10 bases upstream of the archaeal promoter elements BRE (B-recognition element recognized by transcription factor B) and TATA-box (bound by the TATA-binding protein, TBP). The BRE and TATA-box motifs align well with promoter motifs typically found upstream [39] of primary transcription start sites in *H. volcanii* (Fig 4A-C). Half of the genes that show the additional CsmR binding motif are functionally linked to chemotaxis, archaellation, or cell shape determination, supporting a direct role for CsmR regulation beyond motility. In contrast, no significantly enriched motifs were detected in the consistently downregulated genes (n: 12). Notably, 70% (35 out of 50) of the strongly upregulated genes (adjusted *P*-value < 0.05, log_2_-fold change > 1) during exponential phase contained the CsmR binding motif upstream of the first gene in either single- or multi-gene transcription units (Fig 4D). This high enrichment underscores a specific regulatory response linked to *csmR* deletion. Most motif-containing genes displayed growth-phase-dependent expression patterns, with some showing decreasing RNA levels during phase transitions, while others gradually increased. However, these trends were largely consistent between wild type and Δ*csmR* strains (Fig 4E). Interestingly, CsmR itself is stably expressed across all conditions, implying that its regulatory activity may be modulated post-transcriptionally or through interactions with co-regulators. Following the identification of the CsmR binding motif upstream of key archaellation and chemotaxis genes, we sought to further investigate CsmR’s structural classification and its potential DNA-binding properties. Given its proposed role as a transcriptional regulator, we conducted homology modeling and electrostatic surface analysis to determine whether CsmR’s structure supports direct DNA interaction. Foldseek comparisons revealed a strong structural similarity to the Phr regulator from *P. furiosus* (PDB ID: 2P4W), particularly in its dimeric form and predicted DNA-binding interface (S4 Fig). Electrostatic surface mapping identified a positively charged DNA-binding region, closely resembling known DNA-binding sites in Phr regulators, further supporting CsmR’s function as a transcriptional regulator (S5 Fig). Multiple sequence alignment (MSA) across haloarchaeal species demonstrated strong conservation of key basic residues (arginine and lysine) within the predicted DNA-binding domain, reinforcing its potential role in direct transcriptional regulation (S6 Fig). Further analysis using the Protein Structure Transformer (PeSTo) tool [40] supports DNA-binding of CsmR (S7 Fig) predicting a DNA binding interface in the highly conserved and positively charged region. In contrast, the C-terminal region exhibited high sequence variability, consistent with its predicted unstructured nature and a possible role in protein-protein interactions or regulatory modulation. Indeed, the PeSTo-analysis predicts a protein interaction surface at the C-terminus of CsmR, providing a potential interaction site for further regulatory elements (S7 Fig).

**Fig 4.**
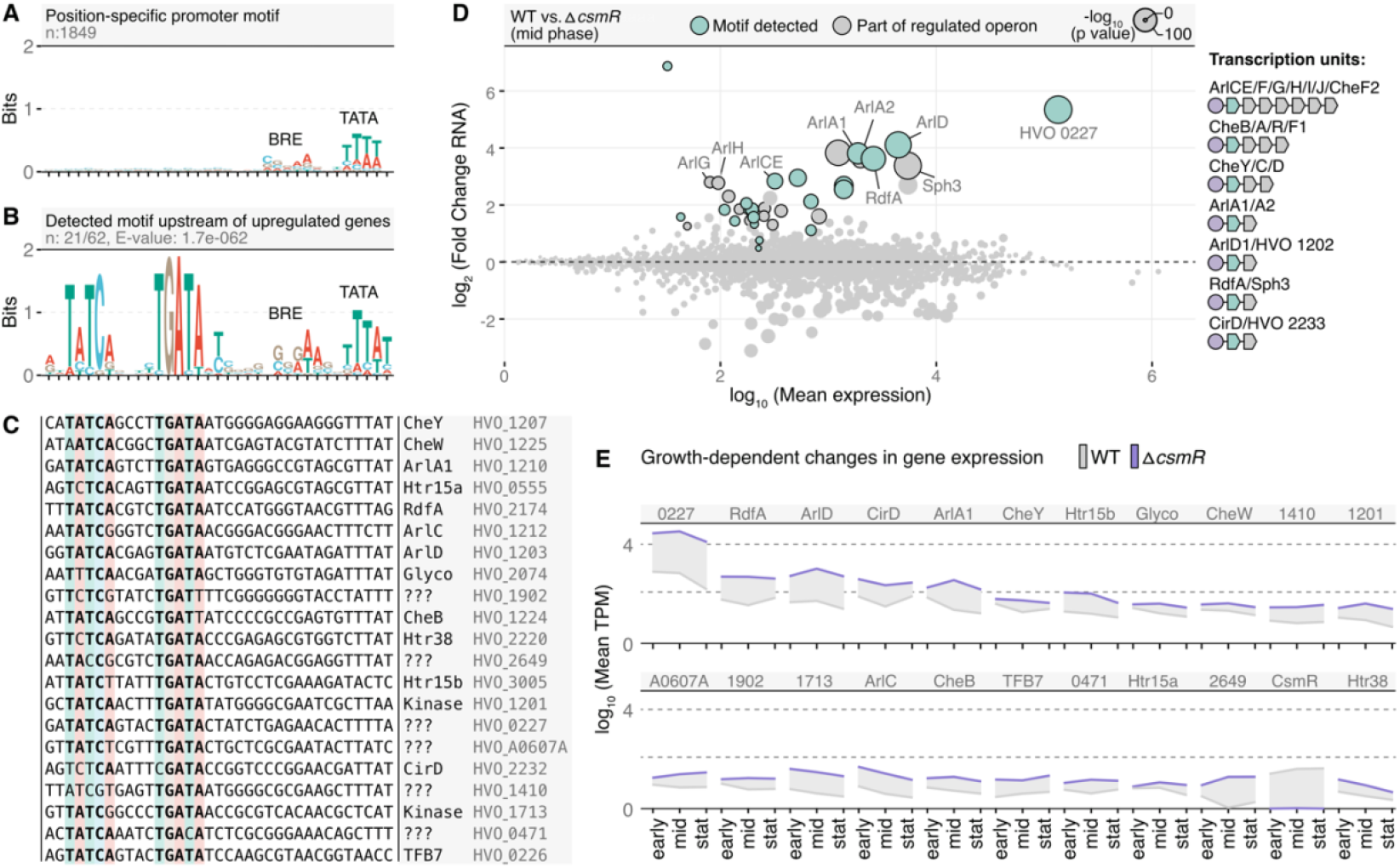
Motif detection and analysis of gene regulation linked to the deletion of *csmR*. (A) Position-specific promoter motif identified in upstream sequences of primary transcription start sites in *H. volcanii*. Archaeal-specific promoter elements, BRE (TB-recognition element recognized by TFB) and TATA-box (TBP-binding element), are highlighted for reference. (B) Motif identified by MEME analysis upstream of genes that are upregulated across all growth conditions (adjusted P-value < 0.1, n: 62). (C) Sequence alignment of genes containing the identified motif, sorted by *P-value*. (D) MA plot illustrating differential gene expression in mid-exponential growth. Each gene is represented as a circle, with circle size indicating the significance of regulation. Genes containing the motif are colored light-green, and those that are part of a regulated unit are outlined in black (transcriptional unit details shown on the right with the motif upstream shown as a purple circle). Non-motif containing genes are shown as grey circles. (E) Plot showing growth-dependent changes in gene expression for wild type (grey) and Δ*csmR* (purple) strains. TPM values (y-axis) are compared across all growth conditions (x-axis) for genes with a detected motif and for the transcriptional regulator CsmR.

### CsmR and CirA co-regulate archaellation and chemotaxis through overlapping but distinct pathways

Given that *csmR* deletion resulted in widespread transcriptional upregulation of archaellation and chemotaxis genes, we next sought to identify additional regulatory factors that may interact with or modulate CsmR function. In particular, *cir* genes (*cirA* / *cirD*) emerged as key candidates, given their established role in archaella regulation and the observed transcriptional shifts in Δ*csmR*. To further explore these connections, we examined *cir* gene expression patterns in Δ*csmR*. In the wild type, transcript levels for *cirA* and *cirD* were comparable across growth phases. However, in the Δ*csmR* strain, *cirD* exhibited consistent upregulation across all phases, while *cirA* showed reduced expression in early and stationary phases but remained unchanged during mid-exponential conditions (S8 Fig). To further analyze their roles, we generated Δ*cirD* and Δ*cirA* deletion strains. Phenotypically, Δ*cirD* resembled Δ*csmR*, with non-motile cells and shape dependency, while Δ*cirA* displayed hypermotility as previously described [32] and a persistent rod-like morphology across all growth phases (Fig 5A-C). Additionally, transmission electron microscopy revealed archaellation of the *cirA* deletion strain even at stationary growth phase in contrast to the *cirD* deletion strain that was not archaellated at all (Fig 5D, 5 E). Notably, a double deletion strain of Δ*csmR* Δ*cirA* resulted in a phenotype identical to Δ*csmR* alone, suggesting that CirA acts upstream or in parallel to CsmR in motility regulation (S9A, C, D Fig). With respect to cell shape the double deletion of *csmR cirD* showed no difference to the wild type. However, the Δ*csmR* Δ*cirD* strain showed slight motility (S9B, E, F Fig). This was surprising as both single deletions were fully non-motile. Despite these morphological differences in Δ*cirA*, growth phase remained the primary driver of transcriptomic variance across all strains (Fig 5F, 5 G). However, Δ*cirA* exhibited significantly more deregulated genes during early and mid-exponential growth, whereas Δ*csmR* and Δ*cirD* displayed stronger transcriptomic shifts during stationary phase (S10 Fig). Gene set enrichment analysis revealed that Δ*cirA* and Δ*csmR* share similar functional effects, particularly among upregulated genes during early and mid-exponential phases (S11 Fig). However, in stationary phase, Δ*csmR* more closely resembled Δ*cirD*, except for motility-related genes, which were not enriched in Δ*csmR*. Focusing on potential CsmR targets identified through promoter motif analysis, we found that Δ*cirA* deletion led to strong upregulation on the respective genes, whereas Δ*cirD* had minimal impact (S12 Fig). While overall expression correlations between the strains were moderate, CsmR target genes were among the most differentially expressed genes in both Δ*cirA* and Δ*csmR* strains (Fig 5H). Notably, despite high correlation between Δ*csmR* and Δ*cirD* during stationary phase, chemotaxis and archaellin genes showed distinct regulation (S13A Fig). Using publicly available Δ*rosR* transcriptomic data, we did not observe transcriptional changes associated with archaellation that correlated with the patterns seen for Δ*csmR*, Δ*cirA*, or Δ*cirD* (S13B, C Fig). To validate these findings, we analyzed the data independently of the previously identified promoter motifs and detected 49 genes significantly upregulated in Δ*csmR* and Δ*cirA* across all growth phases (padj < 0.1) (Fig 5I). STRING network analysis revealed a highly interconnected cluster of archaella assembly, chemotaxis, and signaling genes (Fig 5J). This core included well-characterized genes such as ArlA, ArlB, ArlC, and CheF, alongside transcriptional regulators like TFB and CirA. Interestingly, pilin genes were upregulated in Δ*csmR* and Δ*cirA*, suggesting a compensatory shift towards type IV pili-related functions, such as surface adhesion, in response to disrupted archaella regulation. Glycoproteins and kinases were also linked to this network, reinforcing the influence of both CsmR and CirA on chemotaxis and cellular signaling. Notably, CirD-regulated genes were absent, highlighting its distinct and phase-specific role.

**Fig 5.**
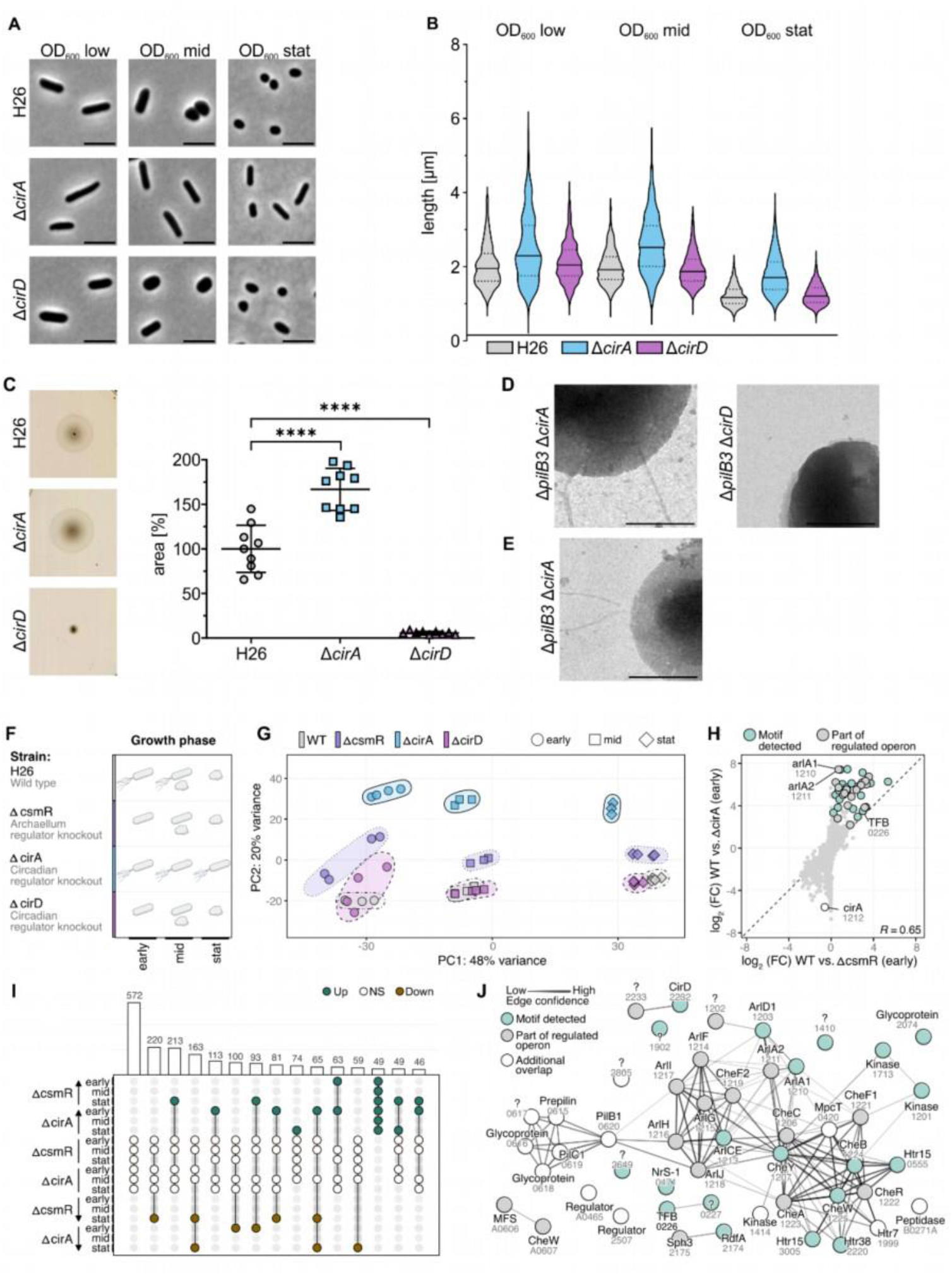
Extensive overlap between CirA and CsmR regulatory networks highlights shared control of archaella-associated genes and motility pathways. (A) Cell shape analysis of the wildtype H26, the *cirA* and *cirD* deletion strain at low, mid and stationary OD_600_. The *cirA* deletion strain stayed rod-shaped over the whole growth cycle and did not transition from rod-to plate shaped cells as the control strain or the *cirD* deletion strain. Scale bar 4 µm. (B) Cell shape was analyzed using MicrobeJ and the summarized results of three independent biological replicates per strain and OD_600_ value plotted as violin-plots. The median is indicated by the middle black line, dotted lines indicate the first and third quartile. For each condition more than 1000 cells were analyzed. (C) Motility assay comparing wildtype H26 with the *cirA* and *cirD* deletion strain. Exemplary motility halos of the three strains are shown. The area of the motility halos was measured and normalized to the average area of wildtype halos showing a significant reduction (p≤0.0001) in cell locomotion of the *cirD* deletion strain compared to H26. in contras the deletion strain of *cirA* showed increased (p≤0.0001) motility compared to the wildtype. Samples were measured in biological and technical triplicates and all nine single data points per strain were plotted. The middle line indicates the mean and the upper and lower line the standard deviation. (D) Transmission electron microscopy images of the *pilB3 cirA* and the *pilB3 cirD* double deletion strains showing that the *cirA* deletion strain is archaellated while the *cirD* deletion strain is not. Scale bar1 µm. (E) Transmission electron microscopy images of the *pilB3 cirA* double deletion strain at the stationary growth phase being still archaellated. Scale bar 1 µm (F), Morphological changes in wild type (H26), Δ*csmR*, Δ*cirA*, and Δ*cirD* strains during early, mid, and stationary growth phases. Wild type cells transition between motile rod-shaped and sessile disk-like morphologies, while Δ*csmR* and Δ*cirD* exhibit growth phase-dependent shape changes, and Δ*cirA* remains rod-shaped across all phases. (G), Principal component analysis (PCA) of transcriptomic data from wild type and deletion strains, showing variance explained by principal components 1 (PC1; 48%) and 2 (PC2; 20%). Biological replicates are color-coded by strain and growth phase (circles: early, squares: mid, diamonds: stationary). (H), Pairwise comparison of fold changes in WT vs. Δ*cirA* and WT vs. Δ*csmR*. Genes with upstream promoter motifs (green circle) and operon membership (grey circle) are highlighted. (I), UpSet plot illustrating the overlap of differentially regulated genes (padj < 0.1) across strains and growth phases. (J), STRING network analysis of overlapping genes, including archaella and chemotaxis components, revealing a highly interconnected regulatory network. High-confidence edges (confidence indicated by line width) indicate functional associations.

### Posttranscriptional regulation of *cirA* by the small RNA *hvo_1211s*

Since CirA appears to be prominently involved in archaellum and cell shape regulation we had a closer CsmR links regulation of motility and cell shape in *Haloferax volcanii* look at its genetic neighborhood. Very recently, the transcriptional regulator RosR was shown by ChIP-seq to bind to the 3’ end of the *arlA2* gene that is located downstream of *cirA* [41]. Interestingly, the cells in the Δ*rosR* mutant were also non-motile ([41], S14A,B Fig) most likely due to the disability to form rod-shaped cells (S14C, D, E Fig) that is essential for *H. volcanii* to swim. Northern blot analysis of RNA isolated from the *rosR* deletion strain showed enriched *cirA* RNA levels compared to the wildtype (S15A, B). This might explain the disability of Δ*rosR* cells to form rod shaped cells that are motile since CirA is involved in the formation of plate-shaped cells. Analysis of transcription start site data at OD_600_ of 0.8 under standard growth conditions [39]revealed a transcription start site downstream of the 3’ end of *arlA2* and the RosR binding site. Northern blot analysis confirmed the expression of an RNA with a size of about 420 nt (S15C Fig). We term this hitherto unannotated gene *hvo_1211s*. With its 420 nt the gene reaches from the intergenic region between *arlA2* and *cirA* into the 3’ end of *cirA* (S16 Fig). Since the *hvo_1211s* RNA is complementary to almost half of the *cirA* RNA it has the potential to act as an anti-sense RNA to regulate *cirA* translation or mRNA stability. To test this hypothesis, we deleted the part of the *hvo_1211s* RNA in the intergenic region leaving *cirA* intact. Similar to Δ*cirA*, partial deletion of *hvo_1211s* showed a hypermotile phenotype compared to the wild type (S17A, B Fig). However, unlike in the *cirA* deletion strain cell shape transition during growth was not affected. Overexpression of the full length *hvo_1211s* blocked rod formation during early growth and hampered motility in the wildtype and partially in the *hvo_1211s* deletion strain (S17 Fig), similar to the *rosR* deletion strain (S14 Fig). The similar phenotype that is caused by *rosR* deletion and *hvo_1211s* overexpression implies an inhibitory function of RosR on *hvo_1211s* transcription that in turn might have a promoting effect on *cirA* transcript abundancy as detected by Northern blotting (S15B, C Fig). Additionally, to exclude that the hypermotility phenotype of Δ*cirA* is a result of the partial deletion of *hvo_1211s*, we generated a partial deletion of *cirA* leaving *hvo_1211s* full sequence intact. The partial *cirA* deletion showed the same phenotype as the full *cirA* knock-out. Cells were rod-shaped over the whole growth cycle and hypermotile (S18A-E Fig). This implies that the cause for the hypermotility of Δ*cirA* is its constant rod-shaped cell shape whilst the cause of the hypermotility of Δ*hvo_1211s* might be a direct impact on the archaellum operon since the deletion strain kept its cell shape like the wild type. Taken together, it is possible that *hvo_1211s* might act as a regulatory sRNA that impacts archaellum genes thereby adding an additional layer of post-transcriptional regulation on cell shape and motility.

## Discussion

### Summary

This study identifies CsmR as a novel transcriptional regulator integrating motility and cell shape control in *H. volcanii*. Our findings establish CsmR as essential for archaellation, linking its function to both the transcriptional regulation of motility genes and the morphological transitions during growth. Unlike previously described archaellum regulators, which typically activate archaellation under favorable conditions [24,25], CsmR deletion leads to a paradoxical upregulation of archaellum and chemotaxis genes, despite the complete loss of motility. This suggests that CsmR is not a simple activator but may act within a more complex regulatory network, including CirA, RosR and *hvo_1211s*, that are involved in the coordination of archaellation, cell shape transitions, and subsequently environmental adaptation.

### Transcriptional control and evolutionary context

A defining feature of *H. volcanii* archaellum regulation is its dispersed operon organization, where motility and chemotaxis genes are interspersed and transcribed in opposing directions [10]. This contrasts with the clustered archaellum loci found in most Euryarchaeota, likely necessitating a complex regulatory framework. The extensive overlap of CsmR and CirA regulons supports a model in which CsmR acts as a central regulator, while CirA fine-tunes motility responses. The deletion of *cirA* results in hypermotility and a prolonged rod morphology [36, our data], whereas *csmR* deletion eliminates motility despite transcriptional upregulation of archaellum genes. The *csmR*/*cirA* double deletion has the same phenotype as the Δ*csmR* mutant, reinforcing the hypothesis that CsmR occupies a dominant regulatory position, with CirA acting as a modulator rather than an independent transcriptional activator.

These findings parallel the multi-layered regulatory strategies observed in Thermoproteota, such as *S. acidocaldarius*, where motility control involves both transcriptional and post-translational mechanisms. In *S. acidocaldarius*, archaellation is regulated by ArnA and ArnB, which are modulated by phosphorylation, enabling rapid environmental responsiveness [14,16]. CirA may exert post-translational control over CsmR function, potentially via the unstructured C-terminal CsmR links regulation of motility and cell shape in *Haloferax volcanii* part, reminiscent of phosphorylation-based repression in *Sulfolobus*. Additionally, the structural similarity of CsmR to Lrp/AsnC-type regulators-widely conserved in archaea and bacteria-suggests evolutionary parallels with bacterial motility regulators, which integrate transcriptional control with metabolic adaptations [42,43]. This is supported by a recent study already linking transcriptional regulation to metabolic control in *H volcanii*, revealing an interplay between cellular development and central metabolism. In this study TrmB and TbsP were identified as a key transcriptional regulators that influence motility and biofilm formation by controlling the glucose metabolism through co-regulation of *gapII* expression [44]. Furthermore, bacterial motility control involves both transcriptional and post-transcriptional regulation through flagellar-specific sigma factors, small RNAs, and kinases to ensuring precise control over flagellum assembly, sometimes even morphology-dependent [45–50]. The identification of *hvo_1211s* includes a regulatory RNA in the archaellum and cell shape regulatory network of *H. volcanii*. Through its overlap with *cirA* initially thought to act as an anti-sense RNA for *cirA* blocking its translation our data suggest that it rather supports *cirA* expression resulting in increased levels of CirA that lead to the formation of plate shaped cells and the reduction of motility that is tightly coupled to cell shape. Thereby *hvo_1211s* might shield the *cirA* mRNA from endonucleolytic cleavage stabilizing the 3’ end of its target by interaction with their complementary parts supporting mRNA stability, similar to the observation of a *sRNA* in the archaeon *Methanosarcina mazei* [51].

The link between motility and cell shape regulation *in H. volcanii* further emphasizes the importance of CsmR. The discovery of key shape-determining factors in *H. volcanii*, including RdfA and DdfA, further underscores the tight coordination between morphology and motility [31]. The persistent rod morphology of Δ*cirA* cells and the normal shape transitions of Δ*csmR* cells indicate that CirA suppresses CsmR-driven motility and shape-determining pathways. The upregulation of pilin genes in both Δ*csmR* and Δ*cirA* mutants suggests regulatory crosstalk between archaella and type IV pili, potentially allowing cells to compensate for disrupted motility by shifting toward surface-associated behaviors [32,52]. Similar compensatory mechanisms have been observed in bacteria, where flagellar defects trigger alternative motility or adhesion pathways [53]. Additionally, ArlI and ArlJ, key players in archaella regulation, have been implicated in pilus-dependent motility, hinting at deeper connections between archaellation and type IV pilin systems [32]. The ability of ArlI and ArlJ mutations to suppress pilin-related motility defects suggests a compensatory mechanism that fine-tunes appendage function, potentially through regulatory crosstalk with CsmR and CirA. Understanding how these systems interconnect will be critical in defining the full regulatory architecture of *H. volcanii* motility.

### Model: CsmR as a central regulator of archaellation

Our findings suggest that CsmR serves as a central regulatory node within the archaella regulatory hierarchy. The deletion of *csmR* results in a complete loss of functional archaella and swimming ability, despite transcriptional upregulation of archaella- and motility-related genes. Based on the position of the CsmR binding motif upstream of BRE and TATA promoter elements in CsmR-targets, it can be inferred that CsmR acts as a positive regulator under standard early growth conditions, potentially enhancing TFB recruitment to ensure proper transcription of all necessary components for archaellum assembly. This positive regulation is counter-balanced by CirA, which functions as a strong local repressor to fine-tune the system for phase-specific regulation.

Deletion of CirA results in hypermotility and upregulation of overlapping regulons, suggesting that CirA prevents overactivation of motility pathways by CsmR. In the absence of CsmR, CirA expression is reduced, which might derepress its targets further. The Δ*csmR* Δ*cirA* double mutant phenotype, resembling Δ*csmR* alone, supports the idea that CsmR occupies a dominant position in the regulatory hierarchy, while CirA functions as a subordinate regulator. Overexpression of CsmR in this model overrides CirA’s repressive influence, resulting in archaellated constantly rod-shaped strains. *CirA*’s mRNA is influenced by the regulatory RNA *hvo_1211s* stabilizing it and *hvo_1211s* in turn is negatively regulated by RosR binding upstream of its promotor region fine tuning the interplay between cell shape transition and motility (Fig 6).

**Fig 6.**
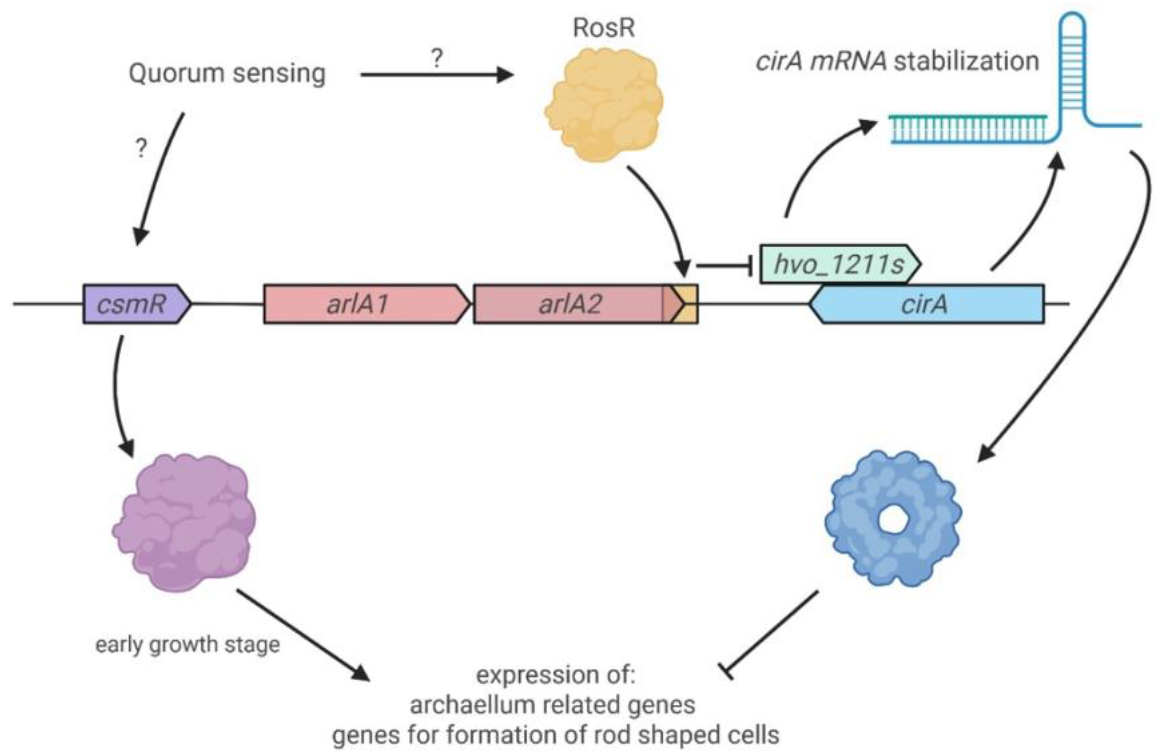
Archaellum and cell shape regulation in *Haloferax volcanii*. CsmR (purple) acts as central regulatory node within the archaellum regulatory network working as positive regulator for cell locomotion during early-stage growth when cells are motile and rod shaped. CirA (blue) counterbalances the positive regulation of CsmR during early growth, fine tuning archaellum expression and rod shape formation. CirA biosynthesis is supported by the RNA transcript *hvo_1211s* (green) stabilizing the *cirA* mRNA by interaction through their homologous regions. *Hvo_1211s* is regulated by RosR (yellow) that binds to the 3’-end of *arlA2* (yellow box) blocking transcription of *hvo_1211s*. CsmR and RosR might be regulated by a quorum sensing signal. The figure was created using BioRender (https://BioRender.com).

This hierarchical model is consistent with growth-phase-specific regulation, that is sensed by a recently described two-component system [54], enabling flexible fine-tuning of activation and repression. It also explains the stage-dependent repression of archaella-related genes, such as Sph3 and RdfA, which contribute to shape-dependent changes. The potential layered regulation by the TFB variant (HVO_0226), CsmR, CirA, *hvo_1211s*, RosR and two additional transcriptional regulators (HVO_A0465, HVO_2507) highlights the complexity of transcriptional control required for precise coordination of cell shape determination motility and archaella assembly in *H. volcanii*.

### Future perspectives and open questions

Our findings establish CsmR as a key transcriptional regulator coordinating motility and cell shape transitions in *H. volcanii*, yet critical questions remain. The identification of a conserved DNA-binding motif suggests direct transcriptional regulation, but its precise mechanism-whether as an activator, repressor, or dual-function regulator-requires final validation. Furthermore, it has to be evaluated if CsmR binds the potential CsmR binding motif in this study and if so, whether it binds as a dimer or monomer. The interplay between CsmR and CirA suggests a layered regulatory system, raising the possibility that CirA modulates CsmR post-translationally, perhaps through phosphorylation. Given that phosphorylation-based archaellum regulation is well-documented in Thermoproteota, a similar mechanism in *H. volcanii* would suggest convergent strategies across archaeal lineages [55]. Beyond transcriptional control, the environmental factors governing CsmR activity remain unclear. Archaellation in *H. volcanii* is influenced by temperature and nutrient availability, suggesting that CsmR integrates external signals to balance motility and morphological transitions [27]. The connection between CsmR and other transcriptional regulators, including alternative TFBs and RosR, should be explored to map its broader regulatory network. Evolutionarily, the structural similarity between CsmR and bacterial Lrp/AsnC regulators highlights potential conserved functions linking motility and metabolism. Moreover, quorum sensing in *H. volcanii*, controlling cell morphology and motility by a small secreted molecule has recently been discovered [56]. This quorum sensing mediated regulation intersects with our described transcriptional network adding another layer of cell shape and motility regulation to its already entangled regulatory network. The upregulation of pilin genes in Δ*csmR* and Δ*cirA* mutants further suggests regulatory crosstalk between archaella and type IV pili systems, pointing to a more integrated control of cell appendages.

By defining CsmR’s role within the archaeal transcriptional landscape, this study provides a foundation for future work on regulatory networks governing motility, environmental sensing, and cellular architecture, with broader implications for the evolution of prokaryotic transcriptional control.

### Material and methods

CsmR links regulation of motility and cell shape in *Haloferax volcanii*

### Growth and media

*Escherichia coli* strains for cloning were grown in LB-medium [57] at 37°C. If necessary, LB-medium was supplemented with 100 µg/ml Ampicillin.

For growth of *Haloferax volcanii* either non-selective YPC-medium or selective Ca-medium [58] was used. For cell shape analysis, CA-medium was supplemented with a trace element solution [29] to improve cell shape homogeneity. Cultures smaller than 5 ml were grown in 15 ml tubes while rotating. Larger cultures for microscopy were grown in 20 ml medium in 100 ml flasks under constant agitation at 120 rpm. Growth temperature for *H*. volcanii was 45°C. Plates were prepared as described before [58]. Motility plates were exactly prepared like described by Patro et al, [59].

Archaeal and bacterial strains used in this study are listed (S1 Table).

### Cloning

For plasmid construction all inserts were amplified using the Phusion®High-Fidelity DNA polymerase obtained from New England Biolabs (NEB) following the manufactures protocol. Plasmids were either assembled via restriction enzyme-based cloning or in vivo ligation [60]. Enzymes for cloning were also obtained from NEB and the manufacturers protocol was followed. For in vivo ligation respective plasmids were linearized via polymerase chain reaction and subsequently cleaned by passing the linearized plasmids through an agarose gel. Approximately 50 ng linearized plasmid was mixed with app. 100 ng insert, the mixture transferred to 50 µl chemically competent *E. coli* and cells were then stored on ice for 30 min. Next, cells were heat shocked at 42°C for 1 min, 450 µl LB-medium was added and the cells incubated for 1.5 h at 37°C under constant shaking. Afterwards the whole cell suspension was plated on LB-plates supplemented with 100 µg/ml Ampicillin and the plates incubated overnight at 37°C. Colonies were transferred to 5 ml LB-medium supplemented with Ampicillin, grown over night and screened via colony PCR for successful plasmid generation the next day. Possible correct assembled plasmids were isolated using the Nucleospin® Plasmid Easy Pure Kit (Macherey-Nagel) and send for sequencing. Used plasmids and primers are listed (S2 Table, S3 Table).

### Transformation in *Haloferax volcanii*

For the genetic modification of *Haloferax volcanii*, cells were transformed using polyethylene glycol [58]. Respective *H. volcanii* strains were grown in 10 ml YPC medium over-night at 45°C under constant rotation. The next day cells were harvested and resuspended in 2 ml buffered spheroplasting solution (1 M NaCl, 27 mM KCl, 50 mM Tris HCl pH 8.5, 15% sucrose (sterile filtered)), transferred to a round bottomed 2 ml reaction tube and centrifuged for 8 min at 3380 g. Supernatant was discarded and the pellet resuspended in 600 µl buffered spheroplasting solution. For one transformation 200 µl of the cell suspension was used. To generate spheroplasts 50 mM ethylenediaminetetraacetic acid (EDTA, pH 8) was added, the tube inverted and incubated for 10 min at room temperature. During incubation 1 µg of demethylated plasmid was mixed with 5 µl EDTA (0.5 M, pH 8) and then filled up to a total volume of 30 µl with unbuffered spheroplasting solution (1 M NaCl, 27 mM KCl, 15% sucrose (sterile filtered)). After 10 min the DNA mixture was added to the tube, mixed by inversion and incubated for 5 min. Subsequently 250 µl 60% polyethylene glycol 600 (diluted in unbuffered spheroplasting solution) was added to the cells and the tube was shaken horizontally for mixing. Thirty minutes later 1.5 ml spheroplast dilution solution (23% salt water, 15% sucrose, 3.75 mM CaCl_2_ (sterile filtered)) was added to the transformants, incubated for 2 min following centrifugation at 3380 g for 8 min. The supernatant was discarded and 1 ml regeneration solution (18% salt water, 1 x YPC-solution, 15% sucrose, 3.75 mM CaCl_2_ (sterile filtered)) added whilst the cell pellet was scratched off the wall. Transformants were then incubated at 45°C for 1.5 h without any agitation. Subsequently, cells were resuspended by tapping the tube and incubated for another 3 h at 45°C under constant rotation. The regenerated transformants were then harvested at 3380 g for 8 min, the supernatant discarded and the cell pellet resuspended in 1 ml transformant dilution solution (18% salt water, 15% sucrose, 3.75 mM CaCl_2_ (sterile filtered)). 100 µl of the resuspended cells were plated on selective Ca-plates and plates incubated in an airtight container at 45°C until colonies were visible.

### Gene deletions in *Haloferax volcanii*

For the generation of deletion strains knock-out plasmids (S2 Table) of the genes to be knocked-out were transformed in the respective background strains as described above. Once colonies were visible after transformation, one colony per strain was transferred into 5 ml of non-selective YPC medium and grown over night at 45°C under constant rotation. The next day the growing culture was diluted back 1:500 into fresh 5 ml YPC-medium and grown over night. This process was repeated to a total of three times. Subsequently three dilutions of the growing culture, starting from 10^-2^, were plated on selective Ca-plates supplemented with 50 µg/ml 5-FOA and 10 µg/ml Uracil. Plates were incubated in an airtight container at 45°C until colonies were visible. To screen possible deletion strains 100 colonies per knock-out attempt were streaked on a YPC-plate and grown for two days at 45°C. Colony PCR was then used to screen for successful deletions using primers indicated in S3 Table.

### Light microscopy

In order to examine the effect of the absence or overexpression of *csmR* on the cell shape of *Haloferax volcanii* cells, phase-contrast microscopy was used. Control strain H26 and the *csmR* deletion strain HTQ289 were either transformed with an “empty” expression plasmid (pTA1392) to complement the auxotrophic marker used in the genetic system or with the *csmR* overexpression plasmid pSVA13922. For microscopy, precultures were inoculated in 5ml Cab-medium and grown overnight at 45°C under constant rotation. The next day cells were diluted back into 20 ml fresh Cab-medium and grown to specific optical densities (OD_600_) under constant shaking at 45°C. When cells reached the desired ODs 5 µl cell culture was spotted on 1% agarose pads dissolved in 18 % salt water. For cell shape analysis Fiji (Version 2.14.0) [61]plug-in MicrobeJ (Version 5.13l) [62]was used. Cells of the *cirA, cirD, rosR, sRNA* deletion strains as well as the double deletion strains of *csmR cirA* and *csmR cirD* were exactly prepared as the *csmR* single deletion strain.

### Motility plates

To assess the effect of the different deletion strains on the ability of cell locomotion semi solid Ca-agar (0.3% w/v) plates were prepared as described above. Control strain H26 and the deletion strains were either transformed with pTA1392 or the indicated plasmids for overexpression of *csmR* or the *sRNA*. One colony of each transformed strain was inoculated in 5 ml Cab-medium and grown over night at 45°C under constant rotation. The next day the OD_600_ of the growing cultures was set to 0.2 by dilution with Ca-medium. Sterile toothpicks were used for inoculation of the motility plates. Plates that harbored strains containing the overexpression plasmid were supplemented with 12 mM Xylose to induce expression. Inoculated plates were airtight packed into plastic bags and incubated at 45°C for 3-4 days. Subsequently plates were scanned and the area of motility halos measured using Fiji. Statistical analysis was done using a paired two-tailed t-test.

### Electron microscopy

Since the *csmR* deletion strain is non motile electron microscopy was used to assess if archaella were present in the mutant strain. To ensure that the observed cell appendages on *Haloferax volcanii* were solely archaella a *pilB3* deletion strain as control and a *pilB3 csmR* double deletion strain was used. Both strains contained plasmid pTA1392 and were grown in 5 ml Ca-medium at 45°C, rotating. The next day the cultures were diluted back into 20 ml Ca-medium and grown to an optical density of 0.01 at 45°C under constant shaking. Cells were harvested and resuspended to a theoretical OD_600_ of 20 in Ca-medium. Next, 5 µl of cells were spotted on a glow discharged carbon coated copper grid (Plano GmbH, Wetzlar, Germany) and incubated for 10 sec. Excess liquid was removed using blotting paper and the cells were stained by addition of 2% uranyl acetate (w/v). Imaging was done with Hitachi HT7800 operated at 100 kV, equipped with an EMSIS Xarosa 20-megapixel CMOS camera. The effect of *cirA* and *cirD* deletion on archaellum formation was also assessed in a *pilB3* deletion background as described for the *csmR* deletion. Since the *cirA* deletion strain showed hypermotility, the *cirA pilB3* deletion strain was also imaged during stationary phase to see if archaella were still assembled.

### Lipid extraction

Lipid were extracted from 3-14 mg freeze-dried Haloferax volcanii cells (variation caused by high salt levels) by applying an acidic Bligh and Dyer protocol, containing 5% trichloroacetic acid (TCA), as described elsewhere [63,64]. In short, freeze-dried cells were resuspended in 2:1:0.8 (v/v) MeOH:CHCl_3_:5% TCA, spiked with 10 µg of the internal standard n-Dodecyl β-maltoside (DDM), vortexed and sonicated for 60 min. Phase separation was obtained by addition of CHCl_3_ and 5% TCA. After centrifugation (2500 x g, 5 min at RT), the bottom phase was collected. To ensure total lipid extraction this procedure was repeated two times more. To remove TCA contaminations, the recovered bottom-phase was washed by adding 1:1:0.9 (v/v) MeOH:CHCl_3_:MQ. After centrifugation (2500 x g, 5 min at RT), the bottom phase was collected, dried under a nitrogen (N_2_) stream, and resuspended in 50 µl MeOH.

### LC-MS analysis of lipids

Samples were analyzed using an Agilent Technologies 1290 Infinity high-performance liquid chromatography (HPLC) system, coupled to a heated electrospray ionization–mass spectrometer (Thermo Q Exactive Plus; Thermo Fisher Scientific). 5 µl was injected into an ACQUITY™ UPLC CSH C18 1.7 µm Column, 2.1 × 150 mm (Waters Chromatography Ireland Ltd) operating at 55 °C with a flow rate of 300 µl/min. Separation of the compounds was achieved by a changing gradient of eluent A (5 mM ammonium formate in water/acetonitrile 40:60, v/v) and eluent B (5 mM ammonium formate in acetonitrile/2-propanol, 10:90, v/v). The following linear gradient was applied: 1) 5% eluent B for 2.5 min, 2) a gradient from 5% to 90% eluent B over 36.5 min, 3) holding for 3 min, 4) returning to 5% eluent B in 0.5 min, 5) and holding for 8 min. The column effluent was injected directly into the Thermo Q Exactive Plus operating in positive and negative ion mode. Settings for positive mode: Spray Voltage: 3.20 |kV|, Capillary temperature: 230 °C, S-lens RF level: 50.0, sheath gas flow: 30, auxiliary gas flow: 5. Settings for negative mode: Spray Voltage: 3.20 |kV|, Capillary temperature: 230 °C, S-lens RF level: 50.0, sheath gas flow: 30, auxiliary gas flow: 20, sweep gas flow: 3, Aux gas heater temperature: 380 °C. Spectral data constituting total ion counts were analyzed using the MacCoss Lab Software: Skyline. The following transition settings were used: Scan range: 133.4-2000 m/z, MS1 filtering: Orbitrap, resolving power 70,000 at m/z 400. MS/MS filtering: DDA, Orbitrap, resolving power: 17,500 at m/z 400. Total ion counts for extracted lipid species (S4 Table) were initially corrected for both the internal standard DDM (m/z 509.30 [M-H]− or m/z 528.34 [M+NH4]+) and the OD (due to high salt concentrations the freeze-dried mass could not be used). To further improve sample comparability, lipid species were normalized for total lipid ion count per sample and subsequently plotted on the y-axis.

### RNA sequencing for differential gene expression analysis

#### RNA extraction

To obtain a comprehensive overview of the effect of the deletion of *csmR, cirA* and *cirD* on the transcriptome, RNA sequencing was used. RNA extraction, library preparation and sequencing has been done as described in [65] with minor modifications. Briefly, RNA was isolated from the deletion strains in lag- (OD_600_ 0.02), exponential- (OD_600_ 0.2), and stationary growth-phase (OD_600_ 2), H26 was used as a control. All strains were transformed with plasmid pTA1392 to avoid the addition of uracil to the selective CA-medium used for growth. To yield enough cells in lag-phase one-liter cultures were grown, cells for the exponential phase in 20 ml and cells in stationary phase in 3 ml. In total per strain and growth phase five replicates were grown. Once cells reached the wanted optical density cells were harvested and resuspended to a theoretical OD_600_ of 5. A maximum of two milliliters of resuspended cells were then transferred two a new reaction tube and pelleted. For RNA isolation the RNeasy® Plus Mini Kit from QIAGEN was used. Pellets were resuspended in 600 µl RLT-plus buffer and RNA was isolated according to the manufacturers protocol. RNA was eluted in 31 µl RNAse free water (Roth®) flash frozen in liquid nitrogen and stored at -80 °C.

#### Library preparation and sequencing

RNA quality was assessed using a Bioanalyzer, and only with an RNA integrity number (RIN) of 8.5 or higher were selected for further processing. Prior to library preparation, RNA was treated with Turbo DNase (Ambion, 1 unit) following the manufacturer’s protocol to eliminate any residual DNA. Four independent biological replicates were prepared for each experimental condition. To remove ribosomal RNA, 2 µg of input RNA was depleted using a ribopool specific to *H. volcanii* (siTOOLs) according to the manufacturer’s instructions.

Library preparation and RNA-sequencing followed the Illumina “Stranded mRNA Prep Ligation” Reference Guide, the Illumina NextSeq 2000 Sequencing System Guide (Illumina, Inc., San Diego, CA, USA), and the KAPA Library Quantification Kit - Illumina/ABI Prism (Roche Sequencing Solutions, Inc., Pleasanton, CA, USA). In brief, omitting the initial mRNA purification step with oligo(dT) magnetic beads, approximately 5 ng of rRNA depleted archaeal RNA was fragmented to an average insert size of 200-400 bases using divalent cations under elevated temperature (94°C for 8 minutes). Next, the cleaved RNA fragments were reverse transcribed into first strand complementary DNA (cDNA) using reverse transcriptase and random hexamer primers. Thereby Actinomycin D was added to allow RNA-dependent synthesis and to improve strand specificity by preventing spurious DNA-dependent synthesis. Blunt-ended second strand cDNA was synthesized using DNA Polymerase I, RNase H and dUTP nucleotides. The incorporation of dUTP, in place of dTTP, quenches the second strand during the later PCR amplification, because the polymerase does not incorporate past this nucleotide. The resulting cDNA fragments were adenylated at the 3’ ends and the pre-index anchors were ligated. Finally, DNA libraries were created using a 15 cycles PCR to selectively amplify the anchor-ligated DNA fragments and to add the unique dual indexing (i7 and I5) adapters. The bead purified libraries were quantified using the KAPA Library Quantification Kit. Equimolar amounts of each library were sequenced on an Illumina NextSeq 2000 instrument controlled by the NextSeq 2000 Control Software (NCS) v1. 5.0.42699, using one 50 cycles P3 Flow Cell with the dual index, single-read (SR) run parameters. Image analysis and base calling were done by the Real Time Analysis Software (RTA) v3.10.30. The resulting .cbcl files were converted into .fastq files with the bcl2fastq v2.20 software. Library preparation and RNA-sequencing were performed at the Genomics Core Facility “KFB - Center of Excellence for Fluorescent Bioanalytics” (University of Regensburg, Regensburg, Germany; www.kfb-regensburg.de).

#### Differential gene expression analysis

Raw sequencing reads in FASTQ format were processed for quality control and adapter trimming using fastp (v0.23.4) with the parameters --cut_front -- cut_tail -q 30, ensuring the removal of low-quality bases. Filtered reads were then aligned to the *Haloferax volcanii* DS2 reference genome (GCF_000025685.1) using Bowtie2 (v2.5.3) with default settings. Sequence alignment files in SAM format were converted to BAM using SAMtools (v1.19.2) for downstream analysis.

Gene-level counts were obtained using feature Counts (part of the RSubread package v2.20.0) with a custom GTF file (gff filtered for entries labeled as gene). Differential expression analysis was performed using DESeq2 (v1.46.0) following Bioconductors guidelines. Principal component analysis (PCA) was applied to variance-stabilized data to assess overall data structure and identify outliers. Replicates identified as outliers (wild type low OD replicate 4, wild type mid OD replicate 1, Δ*csmR* mid OD replicate 4, Δ*cirA* mid OD replicate 3, Δ*cirA* stat OD replicate 1) were removed based on visual inspection. Comparative differential expression analysis was conducted between wild type and deletion strains across different growth conditions to identify growth-dependent transcriptional changes.

#### Functional enrichment analysis based on arCOG classification

To investigate the functional characteristics of differentially expressed genes, we performed a functional enrichment analysis using the Archaeal Clusters of Orthologous Genes (arCOG) classification, as previously described [65,66]. Briefly, arCOGs for *H. volcanii* were retrieved from [67], and gene set enrichment analysis was conducted using the goseq package (v1.58.0) in R. For each growth condition, a background gene set was generated from all detected genes. *P*-values for arCOG term overrepresentation were calculated separately for up- and downregulated genes based on RNA-seq data. Terms were considered significantly enriched at a false discovery rate (FDR) threshold of was determined using a of 0.05.

#### Motif analysis

To identify potential regulatory motifs, sequences 300 bp upstream of the translation start sites of genes upregulated across the three growth condition in the Δ*csmR* strain (adjusted *P*-value < 0.1) were extracted. Motif discovery was performed using MEME (v5.5.6) with parameters -minw 4 -maxw 50. The identified motifs were then imported into R using the memes package (v1.14.0) and visualized with ggseqlogo (v0.2). For comparison, primary transcription start sites were extracted from [39].

#### Additional functional and structural analysis

To explore functional relationships among differentially expressed genes, protein-protein interaction networks were analyzed using the STRING database [68].

Structural predictions were conducted using the AlphaFold Server with AlphaFold 3 model [69]. The predicted structure (CIF file) was downloaded and visualized in ChimeraX [70].

To identify structural homologs, Foldseek [71] was used, revealing 2P4W (DOI: 10.2210/pdb2P4W/pdb) as the closest match [72]. The AlphaFold-predicted structure and 2P4W were aligned using least-squares fitting in ChimeraX (match command).

Electrostatic surface potential was calculated and visualized using Coulombic Surface Coloring in ChimeraX.

To assess sequence conservation, a BLAST search was performed against the non-redundant (nr) database using HVO_1209 as the query sequence. Multiple sequence alignment (MSA) of all hits was generated using Clustal Omega [73] and visualized in ChimeraX, with sequences colored by conservation.

### Northern blot analysis

Total RNA was prepared from *H. volcanii* strain as described before (Nieuwlandt et al 1995). Cells were harvested at exponential phase (OD_650_=0.5) and at stationary phase (OD_650_=1.1-1.4). In addition, total RNA was isolated using the RNeasy Plus Kit (Quiagen) from the following strains: three biological replicates of H26 and Δr*osR*, in each case at an OD_600_ of 0.02 and of 0.2. From all different RNA preparations, 10 µg each were separated on 1.5 % agarose-gel, which were subsequently transferred to a nylon membrane (Biodyne® A, PALL). After transfer, the membrane was hybridized with specific oligonucleotides (primer sequences are listed in S3 Table), which were radioactively labelled with [γ-^32^P]-ATP via polynucleotide kinase treatment to detect the *hvo_1211s, cirA* or *5S* transcript. The signals were recorded by an autoradiography film. For quantification of the *cirA* signal, an additional imaging plate (FujiFilm) was used for recording and the intensities were analysed using the ImageLab Software (Bio-Rad). The *cirA* signal was normalized with the *5S* signal, as a loading control.

## Data deposition

### Data availability

RNA sequencing data are available at the European Nucleotide Archive (ENA, https://www.ebi.ac.uk/ena) under project accession number PRJEB79934 (wild type RNA-seq) and PRJEB86879 (Δ*cirD*, Δ*cirA*, Δ*csmR*).

### Code availability

Documentation and code of all essential analysis steps (tools and custom Rscripts) are available from https://github.com/felixgrunberger/CsmR_Archaellum_Regulation_Hvo.

### Supplementary data

Count data, log2FC data, arCOG data, promoter motif data as excel file available. (S5-S8 Table)

## Authors contributions

PN designed the experiments, supervised FN, AE and KC, generated the *cirD, rosR* and *hvo_1211s* deletion strain and the *csmR* and *hvo_1211a* overexpression plasmid, performed RNA isolation, and manuscript writing. KV performed sample preparation, rRNA depletion, and quality control for RNA sequencing. FG conducted RNA sequencing, performed data analysis, and contributed to manuscript writing. PN and FN analyzed cell shape and motility of different mutant strains. KC analyzed cell shape and motility of *cirA/D* deletion strains and the *csmR cirA* and *csmR cirD* double deletion strains. AE generated the *csmR* and *cirA* deletion mutants, as well as the *csmR cirA* and *csmR pilB3* double deletion strains. SS performed electron microscopy. AS performed Northern-Blot experiments. ME isolated lipids and analyzed their composition. WH grew cells for the lipid analysis. AM and DG acquired funding. SVA conceived the initial idea, contributed to manuscript writing, and acquired funding.

## Acknowledgements

PN was funded by a Momentum grant from the VW Foundation (AZ 94993) awarded to SVA. SS was supported by the SFB1381 by the DFG (German Research Foundation) under project no. 403222701-SFB1381. KC received a stipend from the DAAD (German Academic Exchange Service). SG was funded by the DIVA project (Project number 505545313), funded by the DFG (German Research Foundation). This study was funded by the Deutsche Forschungsgemeinschaft (DFG, German Research Foundation) under Germany’s Excellence Strategy (CIBSS – EXC-2189 – Project ID 390939984). We would like to thank the EM facility at the Faculty of Biology, University of Freiburg, for access to the TEM for generation of data. The TEM (Hitachi HT7800) was funded by the DFG grant (project number 426849454) and is operated by the University of Freiburg, Faculty of Biology, as a partner unit within the Microscopy and Image Analysis Platform (MIAP) and the Life Imaging Center (LIC), Freiburg. Research in the Grohmann lab was supported by the basic funds of the University of Regensburg. The work carried out in the Marchfelder Laboratory was funded by the University of Ulm. We thank Zhengqun Li for the construction of the *cirA* deletion plasmid.

