## Supplementary File for "Control of motility and cell shape of *Haloferax volcanii* is linked by a transcriptional regulator"

**S1 Table Strains used in this study**

| Strain Name | Background strain | Genotype | Source/reference |
|---|---|---|---|
| ***Haloferax volcanii*** | | | |
| H26 | - | Δ*pyrE2* | [1] |
| H119 | DS70 | ΔpyrE2, ΔtrpA, ΔleuB | [2] |
| HTQ277 | H26 | Δ*pyrE2* Δ*cirA* | This study |
| HTQ289 | H26 | Δ*pyrE2* Δ*csmR* | This study |
| HTQ247 | H26 | Δ*pyrE2* Δ*pilB3* | [3] |
| HTQ292 | HTQ247 | Δ*pyrE2* Δ*pilB3* Δ*csmR* | This study |
| HTQ293 | HTQ289 | Δ*pyrE2* Δ*csmR* Δ*cirA* | This study |
| HTQ296 | H26 | Δ*pyrE2* Δ*cirD* | This study |
| HTQ817 | HTQ277 | Δ*pyrE2* Δ*pilB3* Δ*cirA* | This study |
| HTQ820 | HTQ296 | Δ*pyrE2* Δ*pilB3* Δ*cirD* | This study |
| HTQ1005 | HTQ289 | Δ*pyrE2* Δ*csmR* Δ*cirD* | This study |
| HTQ968 | H26 | Δ*pyrE2* partial Δ*hvo_1211s* | This study |
| HTQ1009 | H26 | Δ*pyrE2* partial Δ*cirA* | This study |
| HTQ966 | H26 | Δ*pyrE2* Δ*rosR* | This study |
| ***Escherichia coli*** | | | |
| NEB® 5-alpha | - | *fhuA2Δ(argF-lacZ)U169 phoA glnV44 Φ80Δ(lacZ)M15 gyrA96 recA1 relA1 endA1 thi-1 hsdR17* | New England Biolabs |
| *dam*$^-$/*dcm*$^-$ Competent Cells | - | *ara-14 leuB6 fhuA31 lacY1 tsx78 glnV44 galK2 galT22 mcrA dcm-6 hisG4 rfbD1 R(zgb210::Tn10)* Tet$^S$ *endA1 rspL136* (Str$^R$) *dam13::Tn9* (Cam$^R$) *xylA-5 mtl-1 thi-1 mcrB1 hsdR2* | New England Biolabs |

**S2 Table Plasmids used in this study**

| Plasmids | Description | Primers used | Enzymes used | Source/reference |
|---|---|---|---|---|
| pTA131 | Integrative plasmid with a *pyrE2* selection marker for gene deletions in *H. volcanii* (Amp$^r$) | - | - | [2] |
| pTA1392 | Plasmid for the expression of proteins in *H. volcanii* under control of *p.tnaA* and *pyrE2, hdrB* selection markers. Used to complement uracil auxotrophy (Amp$^r$) | - | - | [4] |
| pSVA5681 | Integrative plasmid for the generation of a *cirA* deletion strain (Amp$^r$) | 11335, 11336; 11337, 11338 | KpnI, XbaI | This study |
| pSVA13742 | Integrative plasmid for the generation of a *csmR* deletion strain (Amp$^r$) | 13704, 13705; 13706, 13707 | in vivo ligation | This study |
| pSVA6082 | Plasmid for the expression of proteins in *H. volcanii* under control of *p.xyl* and *pyrE2, hdrB* selection markers (Amp$^r$) | 13531, 13532 | ApaI, NdeI | This study |
| pSVA13922 | Plasmid for the expression of *csmR* under the control of a xylose promotor (Amp$^r$) | 13828, 14612 | PciI, BamHI (plasmid); NcoI, BamHI (insert) | This study |
| pSVA13767 | Integrative plasmid for the generation of a *cirD* deletion strain (Amp$^r$) | 13780, 13781; 13782, 13783 | in vivo ligation | This study |
| pSVA3999 | Integrative plasmid for the generation of a *pilB3* deletion strain (Amp$^r$) | - | - | [3] |
| pSVA13982 | Integrative plasmid for the generation of a partial *hvo_1211s* deletion strain (Amp$^r$) | 15334, 15335; 15336, 15337 | in vivo ligation | This study |
| pSVA13979 | Integrative plasmid for the generation of a partial *cirA* deletion strain (Amp$^r$) | 15359, 15360; 15361, 15362 | in vivo ligation | This study |
| pSVA13964 | Integrative plasmid for the generation of a *rosR* deletion strain (Amp$^r$) | 15320, 15321; 15322, 15323 | in vivo ligation | This study |

| | | | | |
|---|---|---|---|---|
| pSVA13977 | Plasmid for the expression of *hvo_1211s* under the control of a xylose promotor (Amp$^r$) | 15340, 15363 | NdeI, BamHI | This study |

**S3 Table: Primers used in this study**

| Primer Number/Name | Sequence 5'→3' | Description |
|---|---|---|
| 11066 | TCTAGAGCGGCCGCCAC | Forward primer for the linearization of pTA131 |
| 11067 | GGTACCCAATTCGCCCTATAGTG | Reverse primer for the linearization of pTA131 |
| 11335 | CGGGGTACCTCTGCGAGTTCCTCGCT GTTGTCGTCAC | Forward primer for the amplification of the up-stream region of *hvo_1212* (*cirA*) with KpnI restriction site |
| 11336 | GGTTGTGTTGAAACGGGTGTTGGCT GTCGG | Reverse primer for the amplification of the up-stream region of *hvo_1212* (*cirA*) |
| 11337 | ACCCGTTTCAACACAACCCGGGTCG GTCCTGAAGGCCGGTGGCCCCGATC | Forward primer for the amplification of the down-stream region of *hvo_1212* (*cirA*) with 15 bp overhang to the *cirA* upstream fragment |
| 11338 | TGCTCTAGATCGGCGTCCTCAACAGC ATGAACGACC | Reverse primer for the amplification of the down-stream region of *hvo_1212* (*cirA*) with XbaI restriction site |
| 11339 | GGGTCGAGATACTCTTCGTCCTCGGC TTCC | Forward primer for screening and sequencing of *cirA* deletion and the partial *cirA* deletion strain |
| 11340 | GCGGTGGACATGAACGAGACCTCGA TTAAG | Reverse primer for screening and sequencing of *cirA* deletion and the partial *cirA* deletion strain |
| 13531 | TTCGGGCCCCCCGTCTTTCGTCTCCC GTCTCC | Forward primer for the amplification of a 1 kb up-stream fragment of *hvo_B0025* with an ApaI restriction site |
| 13532 | GAATTCGCATATGCAGTATCCTCATT ACCAGCG | Reverse primer for the amplification of a 1 kb up-stream fragment of *hvo_B0025* with an NdeI restriction site |
| 13704 | GGCGAATTGGGTACCGAGCGCGGT GGTGAC | Forward primer for the amplification of the up-stream region of *hvo_1209* (*csmR*) with 15 complementary bases to linearized pTA131 |
| 13705 | CGGAGTCGGAGCGGTAGATGGAGA TTTATCAGGACG | Reverse primer for the amplification of the up-stream region of *hvo_1209* (*csmR*) with 15 bp overhang to the *csmR* downstream fragment |
| 13706 | GATAAATCTCCATCTACCGCTCCGAC TCCG | Forward primer for the amplification of the down-stream region of *hvo_1209* (*csmR*) with 15 bp overhang to the *csmR* upstream fragment |
| 13707 | GGCGGCCGCTCTAGAAGGTACTTGA CCGTCGTATCTTC | Reverse primer for the amplification of the down-stream region of *hvo_1209* (*csmR*) with 15 complementary bases to linearized pTA131 |
| 13708 | ATAGGTAACCCGCACCTCCG | Forward primer for screening and sequencing of *csmR* deletion strain |

| | | |
|---|---|---|
| 13709 | ACGGTGTCGTTGTAGGTGAG | Reverse primer for screening and sequencing of *csmR* deletion strain |
| 13828 | CATGCCATGGAGTCTGACAGGACCC TCTC | Forward primer for the amplification of *csmR* with NcoI restriction site (cloned in pSVA6082) |
| 14612 | CTAGCTAGCTCAGTCCTGCGTAATCT TCC | Reverse primer for the amplification of *csmR* with BamHI restriction site (cloned in pSVA6082) |
| 13780 | GGCGAATTGGGTACCTTGCGGCCTT CGACGAACTC | Forward primer for the amplification of the up-stream region of *hvo_2232* (*cirD*) with 15 complementary bases to linearized pTA131 |
| 13781 | GGTCGGACACTCGCCCGCC | Reverse primer for the amplification of the up-stream region of *hvo_2232* (*cirD*) |
| 13782 | CGGGCGAGTGTCCGACCGGTAACAA CAATGTACGAAC | Forward primer for the amplification of the down-stream region of *hvo_2232* (*cirD*) with 15 bp overhang to the *cirD* upstream fragment |
| 13783 | GGCGGCCGCTCTAGATGTCGCTCAT GCGTCGAGAG | Reverse primer for the amplification of the down-stream region of *hvo_2232* (*cirD*) with 15 complementary bases to linearized pTA131 |
| 13784 | CCATCGACACTTACGACCCC | Forward primer for screening and sequencing of *cirD* deletion strain |
| 13785 | ATAGGCGTCCTCCGAGAGTC | Reverse primer for screening and sequencing of *cirD* deletion strain |
| 8893 | GCCGACGAGAGCGACCTGAC | Forward primer for screening and sequencing of *pilB3* deletion strain |
| 8894 | CGCGTCGCCATCGTCTGGAG | Reverse primer for screening and sequencing of *pilB3* deletion strain |
| 15320 | GGCGAATTGGGTACCTGCGGGATGA AGCCGTAG | Forward primer for the amplification of the up-stream region of *hvo_0730* (*rosR*) with 15 complementary bases to linearized pTA131 |
| 15321 | GCATATGGAAATGTCATCTGCCTATT TAAACTTTGTC | Reverse primer for the amplification of the up-stream region of *hvo_0730* (*rosR*) |
| 15322 | GACATTTCCATATGCTCGGCTGCGTC CACCGCAC | Forward primer for the amplification of the down-stream region of *hvo_0730* (*rosR*) with 15 bp overhang to the *rosR* upstream fragment |
| 15323 | GGCGGCCGCTCTAGATTTCCGAAGC CGAGCGACTG | Reverse primer for the amplification of the down-stream region of *hvo_0730* (*rosR*) with 15 complementary bases to linearized pTA131 |
| 15324 | TCGCGAAGAACTCGTCAATC | Forward primer for screening and sequencing of *hvo_0730* (*rosR*) deletion strain |
| 15325 | TCAAAGAGCGCCGCGTGAAG | Reverse primer for screening and sequencing of *hvo_0730* (*rosR*) deletion strain |

| | | |
|---|---|---|
| 15334 | GGCGAATTGGGTACCAGAGCAGTCC GTGAACAAGG | Forward primer for the amplification of the up-stream region of *hvo_1211s* with 15 complementary bases to linearized pTA131 |
| 15335 | GCCGCTCGGCAGTCGACGTCG | Reverse primer for the amplification of the up-stream region of *hvo_1211s* |
| 15336 | CGACTGCCGAGCGGCAAAACAGGG AGAGCGCAGCG | Forward primer for the amplification of the down-stream region of the intergenic region between *arlA2* and *cirA* with 15 bp overhang to the *hvo_1211s* upstream fragment |
| 15378 | AATGAAAGGAAAGTTTCGGTG | Reverse primer for the amplification of the down-stream region of *hvo_1211s* with 15 complementary bases to linearized pTA131 |
| 15379 | AACTTTCCTTTCATTAAAACAGGGAG AGCGCAGCG | Forward primer for screening and sequencing of the partial *hvo_1211s* deletion strain |
| 15339 | TGGCACGAGTACGTCGAGTC | Reverse primer for screening and sequencing of the partial *hvo_1211s* deletion strain |
| 15340 | GGAATTCCATATGCAGCGCTGTCGTT TGTTCGG | Forward primer for the amplification of *hvo_1211s* with NdeI restriction site (cloned in pSVA6082) |
| 15363 | CGCGGATCCGACCGACATCGGCCTC GAG | Reverse primer for the amplification of *hvo_1211s* with BamHI restriction site (cloned in pSVA6082) |
| 15359 | GGCGAATTGGGTACCGCTGTCGTTT GTTCGGAGTG | Forward primer for the amplification of the up-stream region of *hvo_1212* (*cirA*) with 15 complementary bases to linearized pTA131 |
| 15360 | CCGAGACCGACATCGGCCTCGAG | Reverse primer for the amplification of the up-stream region of *hvo_1212* (*cirA*) |
| 15361 | CGATGTCGGTCTCGGGGTTGTGTTG AAACGGGTG | Forward primer for the amplification of the *hvo_1212* (*cirA*) region where *hvo_1211s* is coded on with 15 bp overhang to the *cirA* upstream fragment |
| 15362 | GGCGGCCGCTCTAGACACCGAATCC GCCGAATTCG | Reverse primer for the amplification of the down-stream region of the *hvo_1212* (*cirA*) region where *hvo_1211s* is coded on with 15 complementary bases to linearized pTA131 |
| Anti- *cirA* | CGTCAGCGAGTCGAGGACGAGCCG GTCGTAGTTGG | Probe against *cirA* for Northern blot analysis |
| Anti - 5S | CGCAGGTGAGCTTAACTTCCGTGTTC GGG | Probe against *5S-rRNA* for Northern blot analysis |
| Anti - *hvo_1211s* | CCAGCTACTGTCTCCTCGCAGTTCGT CTCGGCTCACCGTCC | Probe against *hvo_1211s* for Northern blot analysis |

**S4 Table: Detailed information of annotated lipids, including: lipid type, group, name, molecular formula, adduct type, theoretical m/z, and the mass error of the annotation.**

| Lipid type | Lipid group | Lipid name | Formula | Adduct type | Theoretical m/z | Mass error (ppm) |
|---|---|---|---|---|---|---|
| Phospholipids | PA | PA (40:0) | C43H89O6P | [M-H]- | 731.6324 | 1.1 |
| Phospholipids | PE | PE (40:0) | C45H94O6NP | [M-H]- | 774.6746 | 1.1 |
| Phospholipids | Me-PGP | Me-PGP (40:0) | C47H98O11P2 | [M-H]- | 899.6512 | 0.8 |
| Phospholipids | Me-PGP | Me-PGP (40:1) | C47H96O11P2 | [M-H]- | 897.6355 | 1.0 |
| Phospholipids | Me-PGP | Me-PGP (40:2) | C47H94O11P2 | [M-H]- | 895.6199 | 1.1 |
| Phospholipids | Me-PGP | Me-PGP (40:3) | C47H92O11P2 | [M-H]- | 893.6042 | 0.8 |
| Phospholipids | Me-PGP | Me-PGP (40:4) | C47H90O11P2 | [M-H]- | 891.5886 | 1.2 |
| Phospholipids | Me-PGP | Me-PGP (45:0) | C52H108O11P2 | [M-H]- | 969.7294 | 0.9 |
| Phospholipids | Me-PGP | Me-PGP (45:2) | C52H104O11P2 | [M-H]- | 965.6981 | 1.0 |
| Phospholipids | PG | PG (40) | C46H95O8P | [M-H]- | 805.6692 | 0.8 |
| Phospholipids | PG | PG (40:1) | C46H93O8P | [M-H]- | 803.6535 | 0.9 |
| Phospholipids | PG | PG (40:2) | C46H91O8P | [M-H]- | 801.6379 | 1.1 |
| Phospholipids | PG | PG (40:3) | C46H89O8P | [M-H]- | 799.6222 | 0.7 |
| Phospholipids | PG | PG (40:4) | C46H87O8P | [M-H]- | 797.6066 | 1.0 |
| Phospholipids | PG | PG (45:0) | C51H105O8P | [M-H]- | 875.7474 | 0.9 |
| Phospholipids | PG | PG (45:2) | C51H101O8P | [M-H]- | 871.7161 | 1.1 |
| Cardiolipin | BPG | BPG (80:0) | C89H182O13P2 | [M-H]- | 1520.2983 | 1.0 |
| Cardiolipin | DGD-PA | DGD-PA (80:0) | C98H195O18P | [M-H]- | 1690.4008 | 0.9 |
| Cardiolipin | S-DGD-PA | S-DGD-PA (80:0) | C98H195O21PS | [M-H]- | 1770.3576 | 0.7 |
| Glycolipids | MGD | MGD (40:0) | C49H98O8 | [M+HCOO]- | 859.7244 | 1.1 |
| Glycolipids | DGD | DGD (40:0) | C55H108O13 | [M+HCOO]- | 1021.7772 | 1.1 |
| Glycolipids | S-DGD | S-DGD (40:0) | C55H108O16S | [M-H]- | 1055.7285 | 0.3 |
| Glycolipids | S-DGD | S-DGD (40:1) | C55H106O16S | [M-H]- | 1053.7129 | 1.8 |
| Glycolipids | S-DGD | S-DGD (40:2) | C55H104O16S | [M-H]- | 1051.6972 | 0.7 |
| Glycolipids | S-DGD | S-DGD (40:3) | C55H102O16S | [M-H]- | 1049.6816 | 1.7 |
| Glycolipids | S-DGD | S-DGD (40:4) | C55H100O16S | [M-H]- | 1047.6659 | 0.8 |
| Glycolipids | S-DGD | S-DGD (45:0) | C60H118O16S | [M-H]- | 1125.8068 | 0.5 |
| Glycolipids | 2S-DGD | 2S-DGD (40:0) | C55H108O19S2 | [M-H]- | 1135.6853 | 0.6 |
| Glycolipids | S-Gly-AHH | S-Gly-AHH (40:0) | C55H111O16NS | [M+NH4]+ | 1091.7962 | 0.8 |
| Archaeol | Archaeol | Archaeol (40:0) | C43H88O3 | [M+NH4]+ | 670.7072 | 0.9 |
| Archaeol | Archaeol | Archaeol (40:1) | C43H86O3 | [M+NH4]+ | 668.6915 | 1.1 |
| Archaeol | Archaeol | Archaeol (40:2) | C43H84O3 | [M+NH4]+ | 666.6759 | 0.6 |
| Archaeol | Archaeol | Archaeol (40:3 | C43H82O3 | [M+NH4]+ | 664.6602 | 0.9 |
| Archaeol | Archaeol | Archaeol (40:4) | C43H80O3 | [M+NH4]+ | 662.6446 | 0.7 |
| Archaeol | Archaeol | Archaeol (45:0) | C48H98O3 | [M+NH4]+ | 740.7854 | 1.2 |
| Archaeol | Archaeol | Archaeol (45:2) | C48H94O3 | [M+NH4]+ | 736.7541 | 0.5 |

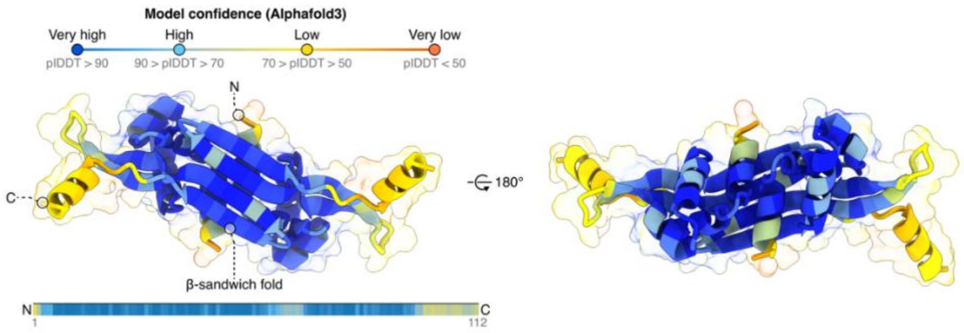

**S1 Fig. AlphaFold3 structural prediction classifies CsmR as an Lrp/AsnC family regulator.**
AlphaFold3 structural prediction of a CsmR dimer, shown in two orientations. The confidence of the model is color-coded from high (blue) to low (yellow/red) based on the pLDDT score (per-residue measure of local confidence). The core structure is defined by a β-sandwich fold, which is highlighted and characteristic of the Lrp/AsnC family of transcriptional regulators. The per-residue confidence score is plotted below the structure.

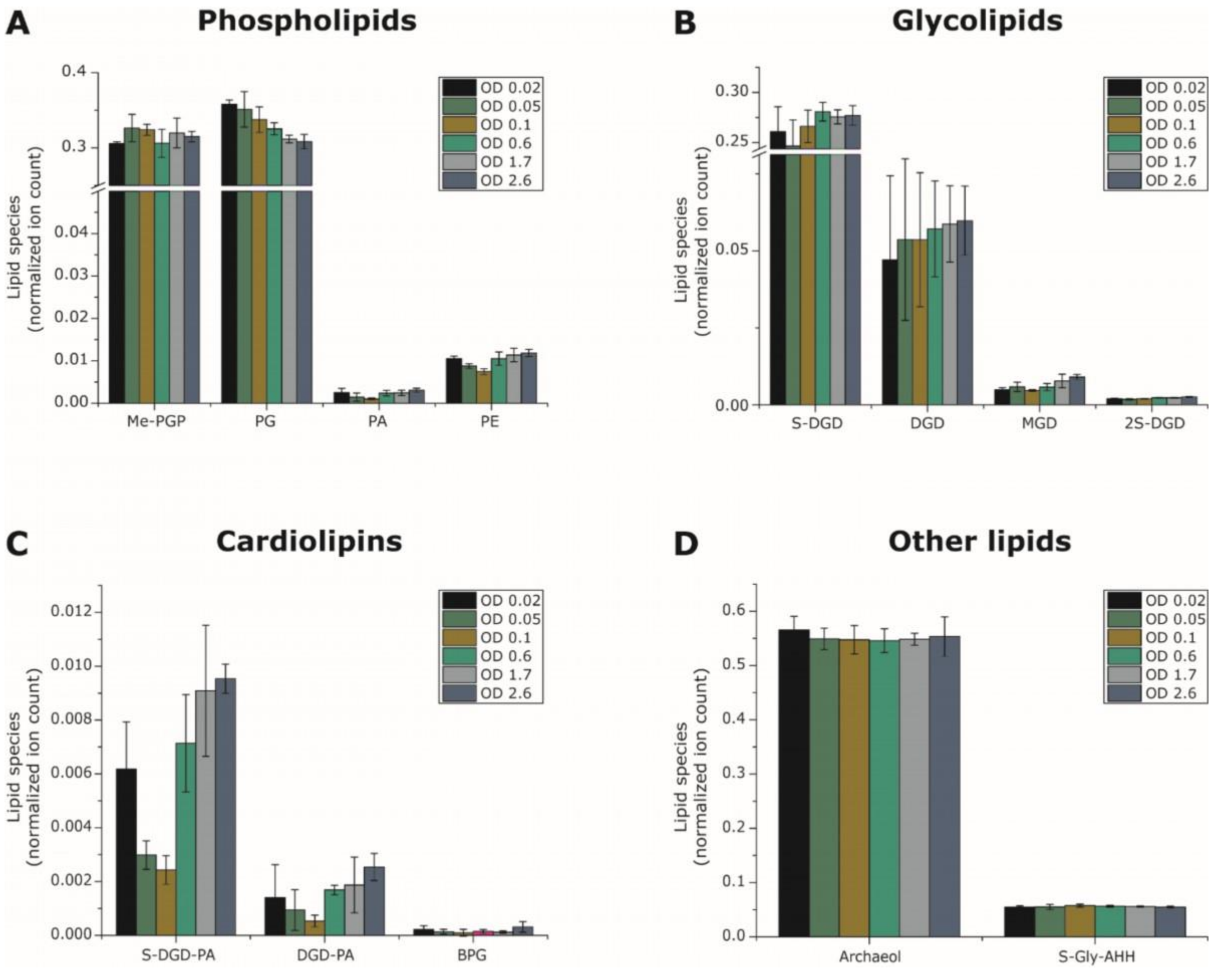

**S2 Fig. Lipid species abundance throughout *H. volcanii* growth stages.**

Lipid species are presented as normalized ion counts. N=4. Presented lipid species: phosphatidylglycerophosphate methyl ester (Me-PGP), phosphatidylglycerol (PG), phosphatidic acid (PA), phosphatidylethanolamine (PE), sulfo-digalactosyldiacylglycerol (S-DGD), digalactosyldiacylglycerol (DGD), monogalactosyldiacylglycerol (MGD), disulfo-digalactosyldiacylglycerol (2S-DGD), BPG (bi-phosphatidylglycerol)/aCL (archaeal cardiolipin), S-Gly-AHH (sulphated glycosylaminohexanehexaol [5,6]).

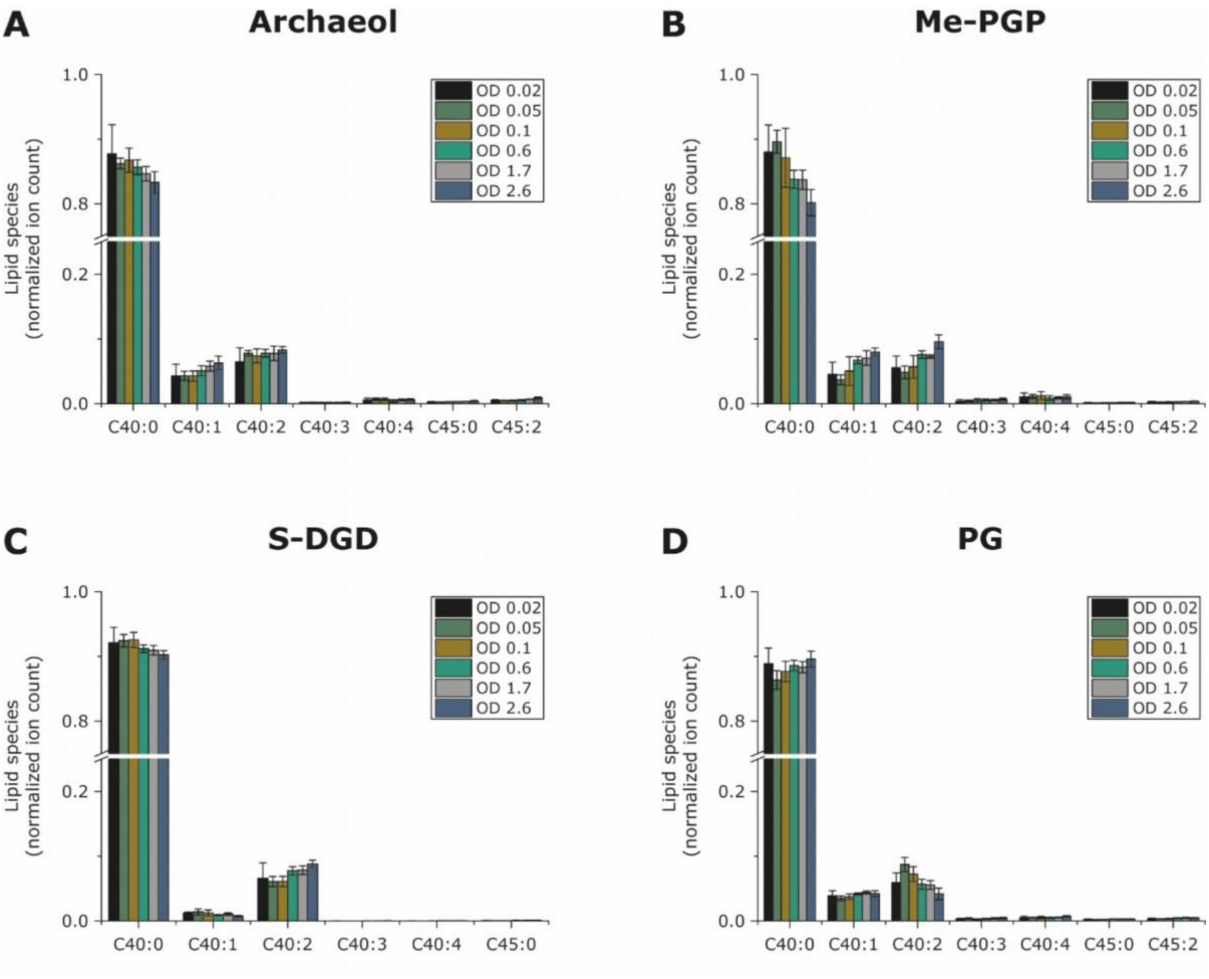

**S3 Fig. Lipid tail distribution of the four most abundant lipid species in *H. volcanii*, during different growth stages.**

(A) archaeol, (B) phosphatidylglycerophosphate methyl ester (Me-PGP), (C) sulfo-digalactosyldiacylglycerol (S-DGD) and (D) phosphatidylglycerol (PG). Lipid species are presented as normalized ion counts. N=4.

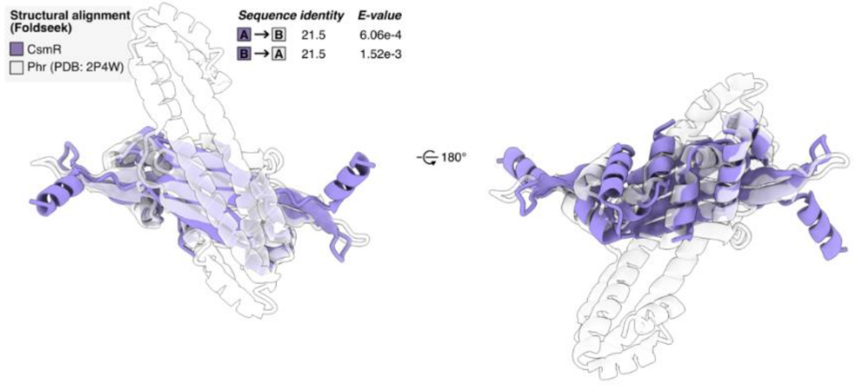

**S4 Fig. Structural alignment of CsmR with the Phr regulator (PDB: 2P4W) using Foldseek.**
CsmR (purple) and Phr (gray) are shown in two orientations to highlight structural similarity. Despite low sequence identity (21.5%), structural alignment reveals a conserved core fold, particularly in the β-sandwich domain, suggesting a shared architectural framework. The E-values indicate statistically significant similarity

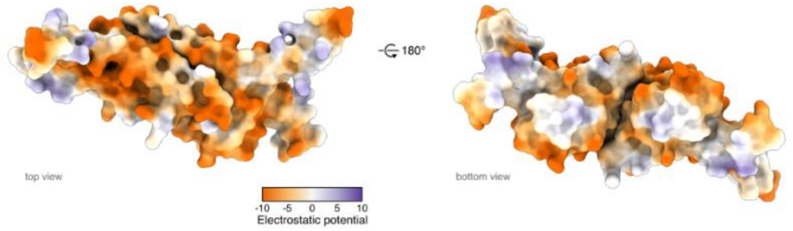

**S5 Fig. Electrostatic surface potential of CsmR.**
Electrostatic surface representation of CsmR shown in two orientations (top and bottom view). The color gradient represents the electrostatic potential, ranging from negatively charged regions (orange) to positively charged regions (purple).

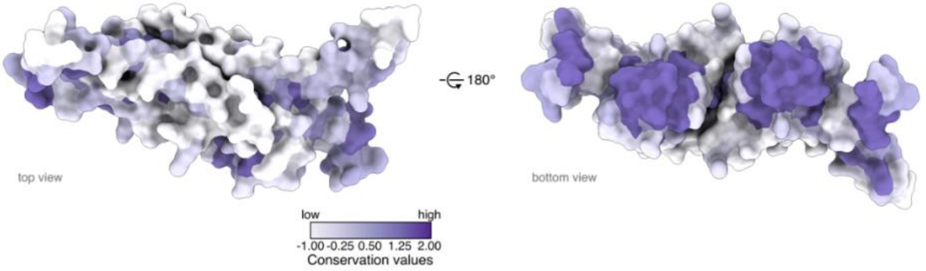

**S6 Fig. Evolutionary conservation of CsmR surface residues.**
Surface representation of CsmR colored by sequence conservation across halobacterial homologs, based on entropy-based conservation scores from AL2CO. Highly conserved residues are shown in dark purple, while variable regions are in white.

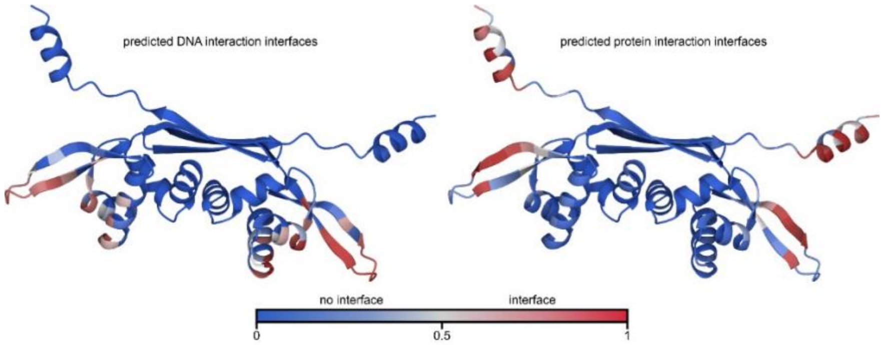

**S7 Fig. DNA and protein interaction interface prediction for CsmR**
DNA-protein and protein-protein interface prediction of the CsmR dimer retrieved from Alphafold3 prediction. Prediction confidence is shown by a color gradient from blue, no interface, to red for interface.

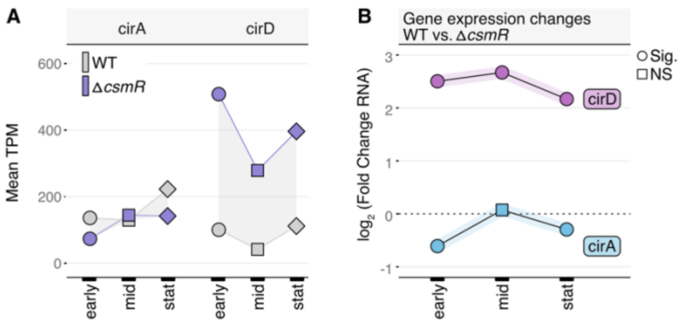

**S8 Fig. Comparative expression analysis of *cirA* and *cirD* genes across growth phases in WT and Δ*csmR* strains.** (A) Mean transcripts per million (TPM) values of *cirA* and *cirD* across early (circle), mid (square), and stationary (diamond) growth phases in WT (grey) and Δ*csmR* (purple) strains. (B) $Log_2$ fold-change in RNA expression (ΔcsmR vs. WT) for *cir* genes across growth phases. Significant genes are depicted as circles, and non-significant genes as squares. Color indicates specific *cir* genes (*cirA*: light blue, *cirD*: purple).

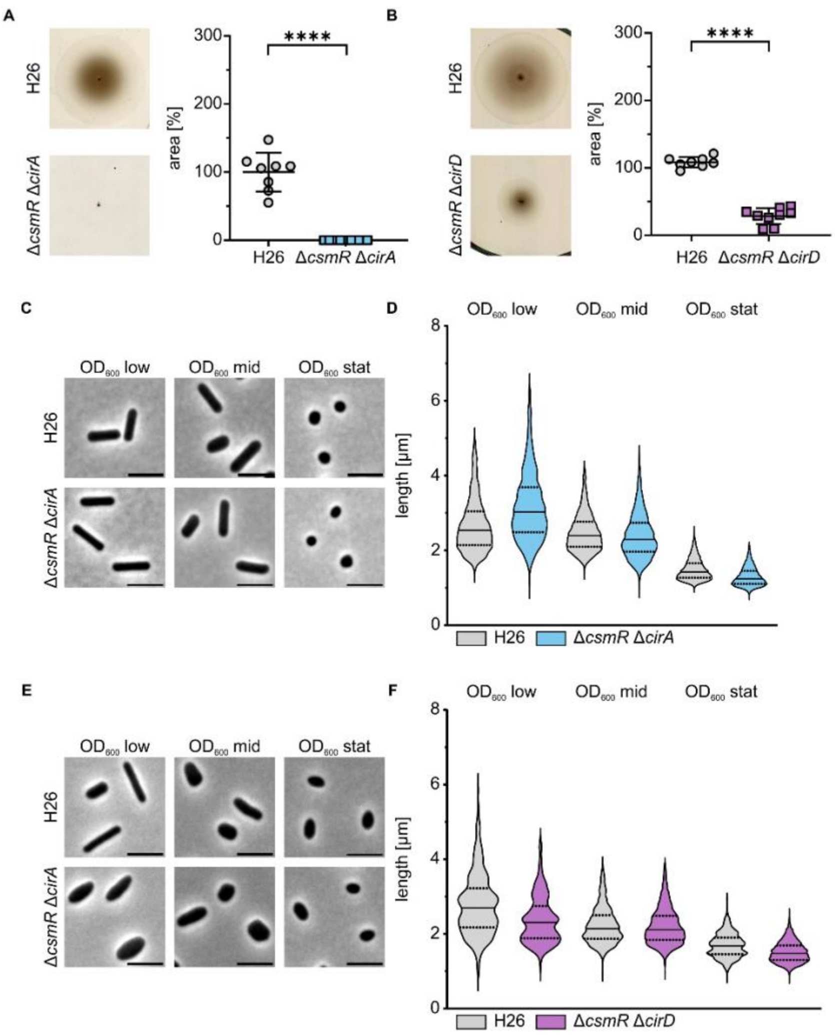

**S9 Fig. Double deletions of *csmR cirA* and *csmR cirD* show similar phenotypes as the *csmR* single deletion mutant.**

(A) Motility assay comparing wild type H26 with the *csmR cirA* double deletion strain. Exemplary motility halos of both strains are shown. The area of the motility halos was measured and normalized to the average area of wild type halos showing significantly ($p \leq 0.0001$) decreased motility in the double deletion strain compared to the H26 control. Samples were measured in biological and technical triplicates and all single data points per strain were plotted. The middle line indicates the mean and the upper and lower line the standard deviation. (B) Motility assay of H26 and the *csmR cirD* double deletion strain. Exemplary motility halos of both strains are shown. The area of the motility halos was measured and normalized to the average area of wild type halos showing significantly ($p \leq 0.0001$) decreased motility in the double deletion strain compared to the H26 control. Samples were measured in biological and technical triplicates and all single data points per strain were plotted. The middle line indicates the mean and the upper and lower line the standard deviation. (C) Cell shape analysis of the wild type H26 and the *csmR cirA* doubled deletion strain at low, mid and stationary $OD_{600}$. Scale bar 4 µm. (D) Cell shape was analyzed using MicrobeJ and the summarized results of three independent biological replicates per strain and $OD_{600}$ value plotted as violin-plots. The median is indicated by the middle black line, dotted lines indicate the first and third quartile. For each condition more than 1000 cells were analyzed. (E) Cell shape analysis of the wild type H26 and the *csmR cirD* doubled deletion strain at low, mid and stationary $OD_{600}$. Scale bar 4 µm. (F) Cell shape was analyzed as described in (D).

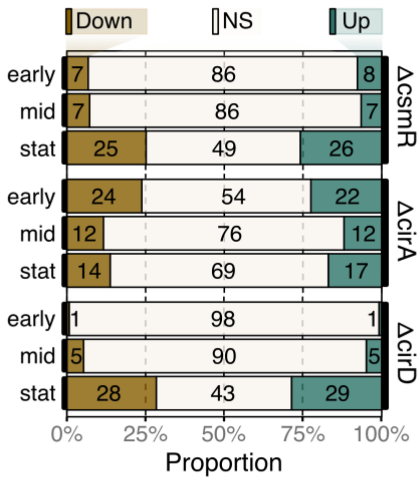

**S10 Fig. Gene regulation across strains and growth phases.**

Proportions of differentially expressed genes (upregulated, downregulated, or non-significant) in Δ*csmR*, Δ*cirA*, and Δ*cirD* strains compared to wild type across early, mid, and stationary growth phases. Bar segments indicate the percentage of genes in each category, with upregulated genes in green, downregulated genes in brown, and non-significant genes in beige

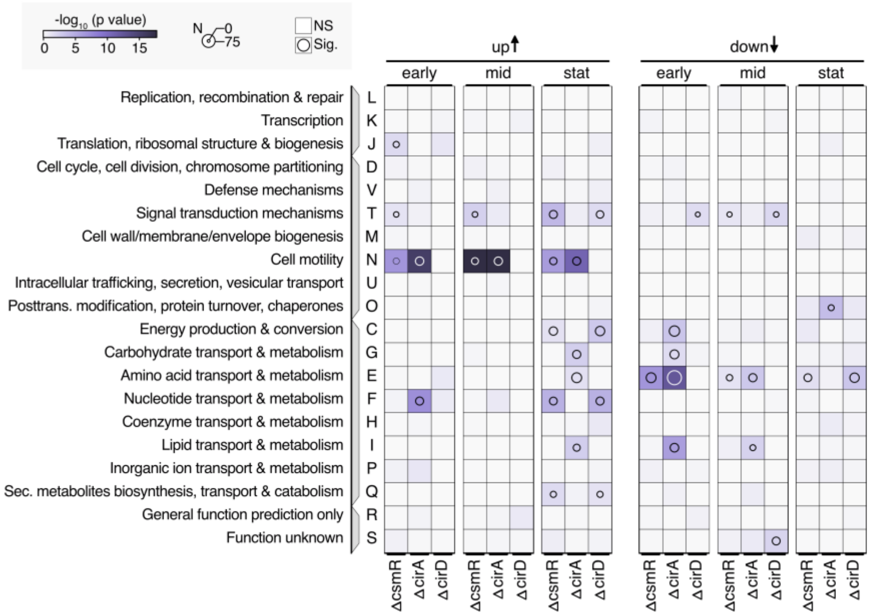

**S11 Fig. Deletion of *cir* genes reveals functional similarities between cirA and csmR in motility regulation, with distinct patterns in stationary phase.**

Gene set enrichment analysis of archaeal clusters of orthologous groups (arCOGs) across all deletion strains and growth phases. Significantly overrepresented genes (P-value < 0.05, color-coded by a white to dark purple gradient) are marked by circles, with circle size representing the total number of differentially regulated genes detected in each group. Separate analyses were performed for upregulated (padj<0.05, log2FC >= 1) and downregulated (padj<0.05, log2FC <= -1) genes across early, mid, and stationary growth conditions.

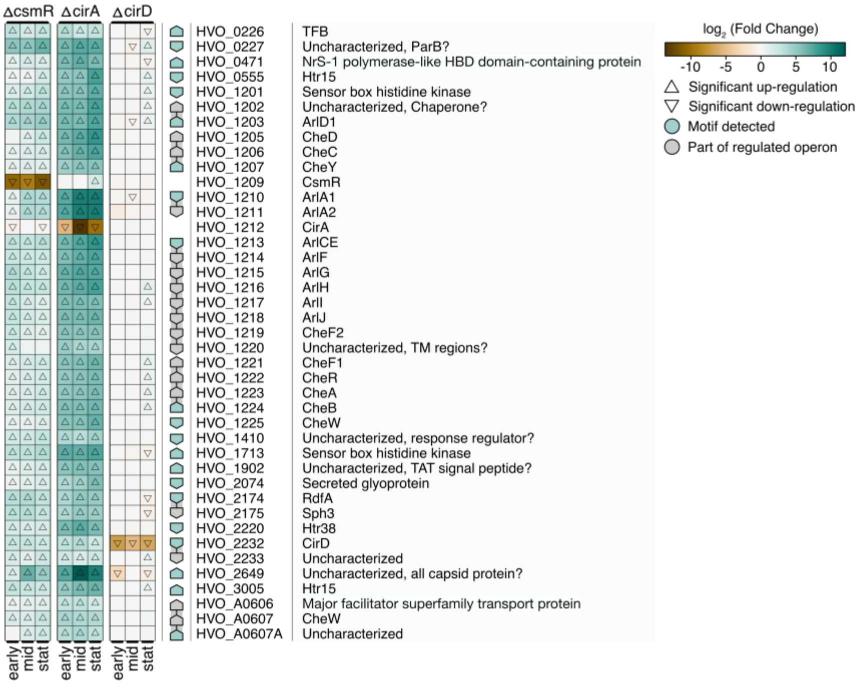

**S12 Fig. Differential expression of potential csmR regulons across deletion strains.**

Log2 fold-change analysis (color-coded from brown to green) of genes in Δ*csmR*, Δ*cirA*, and Δ*cirD* strains across early, mid, and stationary phases. Significant upregulation (up triangle) and downregulation (down triangle) are shown, with motifs detected upstream (light green) and operon membership (grey) indicated.

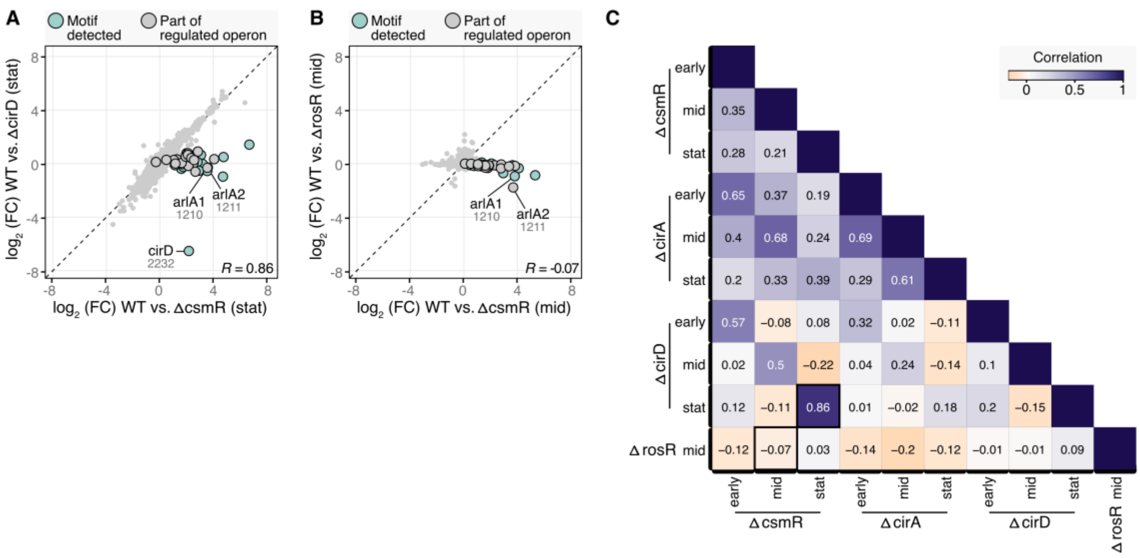

**S13 Fig. Correlation analysis of differential gene expression between deletion strains highlights distinct and shared effects on gene regulation.**

(A) Scatter plot comparing log2 fold changes (WT vs. *ΔcsmR* and WT vs. *ΔcirD*) during the stationary phase and (B) WT vs. Δ*csmR* and WT vs. Δ*rosR* during the mid-phase. Pearsons's correlation was calculated based on pairwise complete observations. Genes with upstream promoter motifs (green circle) and operon membership (grey circle) are highlighted. (C) Heatmap of correlation coefficients for log2 fold changes across strains (Δ*csmR*, Δ*cirA*, Δ*cirD*, Δ*rosR*) and growth phases (early, mid, stationary).

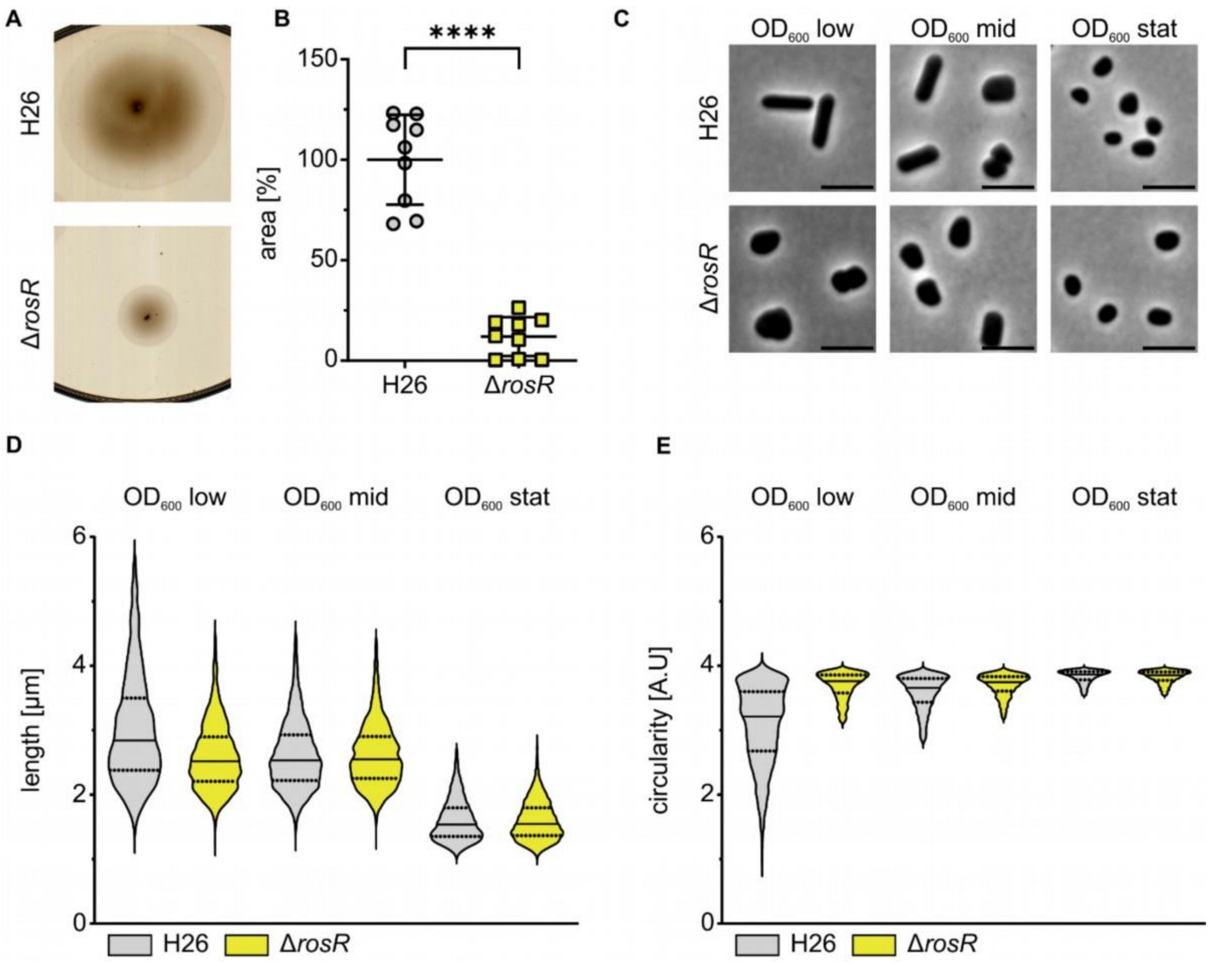

**S14 Fig. Deletion of *rosR* impacts motility and cell shape.**

(A) Motility assay comparing wild type H26 with the *rosR* deletion strain. Exemplary motility halos of both strains are shown. (B) The area of the motility halos was measured and normalized to the average area of wild type halos showing significantly ($p \leq 0.0001$) decreased motility of the *rosR* deletion strain compared to H26. Samples were measured in biological and technical triplicates and all single data points per strain were plotted. The middle line indicates the mean and the upper and lower line the standard deviation. (C) Exemplary images of the wild type H26 and the *rosR* deletion strain at low, mid and stationary $OD_{600}$. Deletion of *rosR* inhibits rod formation. Scale bar 4 µm. (D) Cell shape was analyzed using MicrobeJ and the summarized results of three independent biological replicates per strain and $OD_{600}$ value plotted as violin-plots. The median is indicated by the middle black line, dotted lines indicate the first and third quartile. (E) Cell circularity was analyzed using MicrobeJ and the summarized results of three independent biological replicates per strain and $OD_{600}$ value plotted as violin-plots. The median is indicated by the middle black line, dotted lines indicate the first and third quartile. For each condition more than 1000 cells were analyzed.

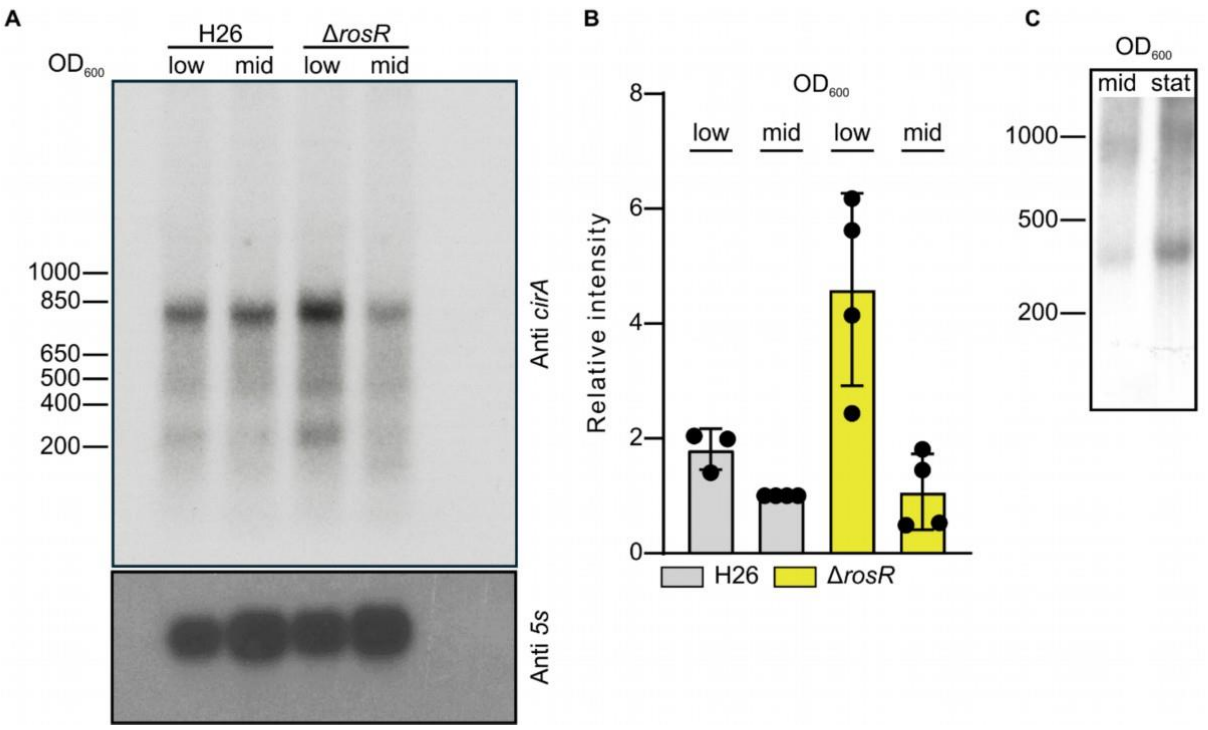

**S15 Fig. Northern blot analysis.**

(A) Northern blot analysis using an anti *cirA* probe on RNA extracted from the wild type H26 and the *rosR* deletion strain at low ($OD_{600}$ 0.02) and mid ($OD_{600}$ 0.2) growth. Anti *5S* probe was used as loading control. (B) Quantification of the signal by the *cirA* probe normalized to the *5S* signal. Plotted are the results from at least three independent experiments per strain and growth phase. (C) Detection of the *hvo_1211s* transcript using RNA extracted from H26 on a Northern blot with a size of approximately 400 nucleotides.

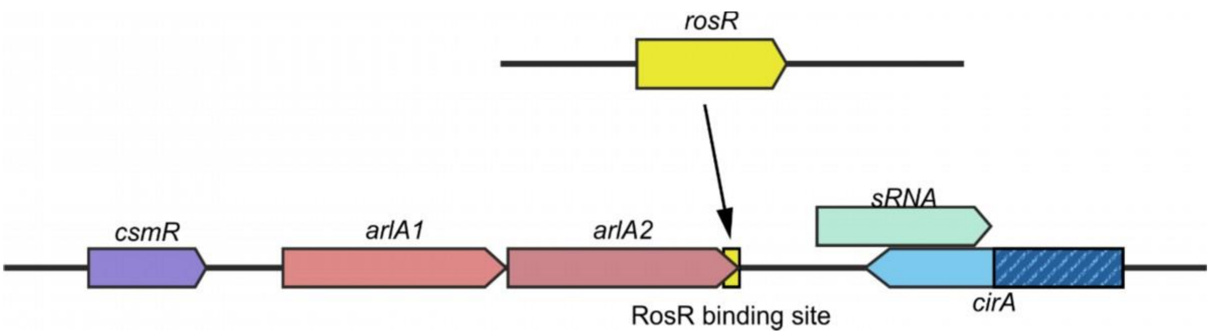

**S16 Fig. Schematic overview of the genetic region between *csmR* (*hvo_1209*) and *cirA* (*hvo_1212*).**
RosR binding to the 3’ end of *arlA2* is indicated by a yellow box. The removed part of the partial *cirA* deletion is indicated in dark blue.

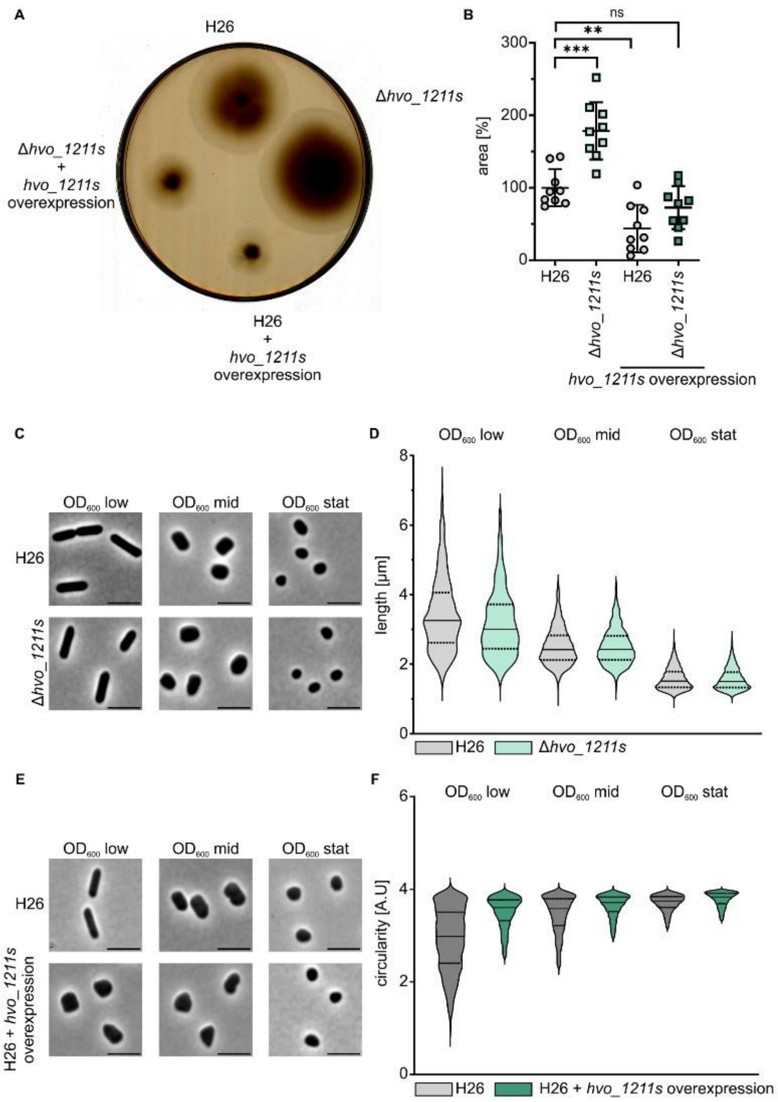

**S17 Fig. Motility of *H. volcanii* is suppressed by a small RNA.**

(A) Motility assay comparing wild type H26 with the *hvo_1211s* deletion strain with and without a plasmid for overexpression of *hvo_1211s*. A exemplary motility plate is shown. (B) The area of the motility halos was measured and normalized to the average area of wild type halos showing significantly ($p \leq 0.05$) increased motility in the *hvo_1211s* deletion strain compared to the H26 control. Overproduktion of *hvo_1211s* suppresses motility in H26 significantly compared to the wild type control ($p \leq 0.05$) and fully complements the hypermotility phenotype of the *Δ hvo_1211s* strain back to wild type levels. Samples were measured in biological and technical triplicates and all single data points per strain were plotted. The middle line indicates the mean and the upper and lower line the standard deviation. (C) Cell shape analysis of the wild type H26 and the *hvo_1211s* deletion strain at low, mid and stationary $OD_{600}$. The *hvo_1211s* deletion strain showed the same phenotype as the control strain. Scale bar 4 µm. (D) Cell shape was analyzed using MicrobeJ and the summarized results of three independent biological replicates per strain and $OD_{600}$ value plotted as violin-plots. The median is indicated by the middle black line, dotted lines indicate the first and third quartile. For each condition more than 1000 cells were analyzed. (E) Cell shape analysis of H26 with and without an *hvo_1211s* overexpression plasmid at low, mid and stationary $OD_{600}$. Overexpression of *hvo_1211s* inhibits rod formation of the cells. Scale bar 4 µm. (F) Cell circularity was analyzed using MicrobeJ and the summarized results of three independent biological replicates per strain and $OD_{600}$ value plotted as violin-plots. The median is indicated by the middle black line, dotted lines indicate the first and third quartile. For each condition more than 1000 cells were analyzed.

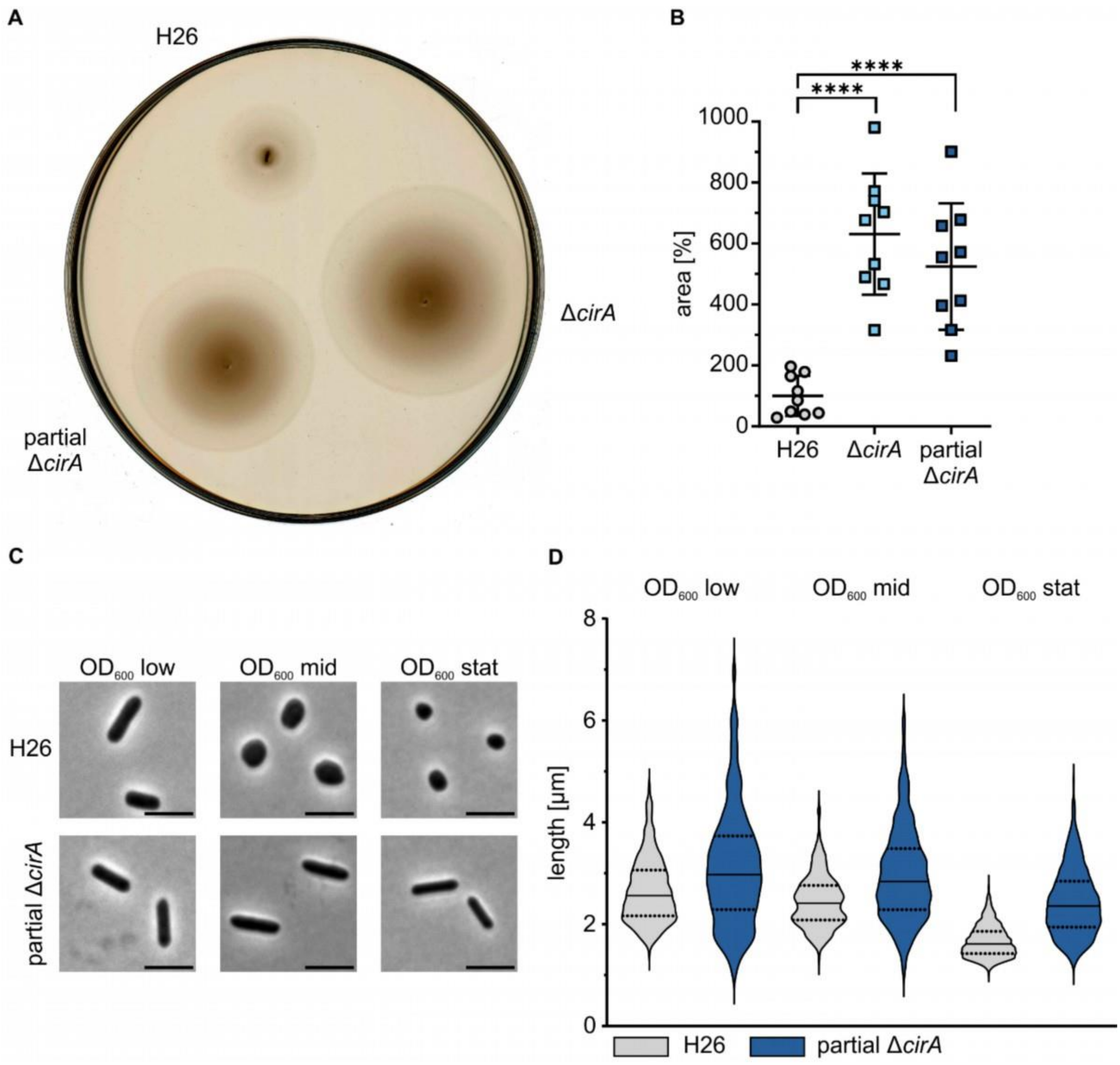

**S18 Fig. A partial deletion of *cirA* leaving the *hvo_1211s* intact shows the same phenotype as Δ*cirA*.**

(A) Motility assay comparing wild type H26 with the *cirA* deletion strain and the partial *cirA* deletion strain. A exemplary motility plate is shown. (B) The area of the motility halos was measured and normalized to the average area of wild type halos showing significantly ($p \leq 0.0001$) increased motility in the *cirA* and the partial *cirA* deletion strain deletion strain compared to the H26 control. Samples were measured in biological and technical triplicates and all single data points per strain were plotted. The middle line indicates the mean and the upper and lower line the standard deviation. (C) Cell shape analysis of the wild type H26 and the partial *cirA* deletion strain at low, mid and stationary $OD_{600}$. The partial *cirA* deletion strain showed the same phenotype as the full Δ*cirA* strain. Cells stayed rod-shaped during the full growth cycle. Scale bar 4 µm. (D) Cell shape was analyzed using MicrobeJ and the summarized results of three independent biological replicates per strain and $OD_{600}$ value plotted as violin-plots. The median is indicated by the middle black line, dotted lines indicate the first and third quartile. For each condition more than 1000 cells were analyzed.
